# EPINEST, an agent-based model to simulate epidemic dynamics in large-scale poultry production and distribution networks

**DOI:** 10.1101/2023.07.25.550458

**Authors:** Francesco Pinotti, José Lourenço, Sunetra Gupta, Suman Das Gupta, Joerg Henning, Damer Blake, Fiona Tomley, Tony Barnett, Dirk Pfeiffer, Md. Ahasanul Hoque, Guillaume Fournié

**Affiliations:** University of Oxford, Oxford, UK; BioISI (Biosystems and Integrative Sciences Institute), Faculdade de Ciências da Universidade de Lisboa, Lisbon, Portugal; School of Veterinary Science, The University of Queensland, Queensland, Australia; Gulbali Institute, Charles Sturt University, Wagga Wagga, NSW, Australia; Royal Veterinary College, London, UK; The Firoz Lalji Centre for Africa, London School of Economics and Political Science, London, UK; City University of Hong Kong, Hong Kong SAR, Hong Kong; Chattogram Veterinary and Animal Sciences University, Chittagong, Bangladesh; INRAE, VetAgro Sup, UMR EPIA, Université de Lyon, Marcy l’Etoile, 69280, France; INRAE, VetAgro Sup, UMR EPIA, Université Clermont Auvergne, Saint Genes Champanelle, 63122, France

## Abstract

The rapid intensification of poultry production raises important concerns about the associated risks of zoonotic infections. Here, we introduce EPINEST (EPI-demic NEtwork Simulation in poultry Transportation systems): an agent-based modelling designed to simulate pathogen transmission within realistic poultry production and distribution networks. The modular structure of the model allows for easy parameterization to suit specific countries and system configurations. Moreover, the framework enables the replication of a wide range of eco-epidemiological scenarios by incorporating diverse pathogen life-history traits, modes of transmission and interactions between multiple strains and/or pathogens. EPINEST was developed in the context of an interdisciplinary multicentre study conducted in Bangladesh, India, Vietnam and Sri Lanka, and will facilitate the investigation of the spreading patterns of various health hazards such as avian influenza, *Campylobacter, Salmonella* and antimicrobial resistance in these countries. Furthermore, this modelling framework holds potential for broader application in veterinary epidemiology and One Health research, extending its relevance beyond poultry to encompass other livestock species and disease systems.

## Introduction

Animal populations act as reservoirs for a wide range of zoonotic pathogens, such as Ebola virus, MERS-CoV, SARS-CoV-2, avian influenza viruses (AIVs), *Campylobacter* and *Salmonella* [1–6]. Within this, livestock production is known to promote the risk of zoonotic infections [7]. In the case of emerging pathogens of wildlife, livestock may become intermediate or amplifier hosts, increasing odds of spillover into the human population [8]. The ongoing global intensification of livestock production raises critical questions about the role of husbandry and animal trading practices in shaping the risk of zoonotic epidemics or spillover events. [9, 10]. Unfortunately, however, a comprehensive understanding of how suck risk is modulated and amplified along production and distribution networks (PDNs) is lacking.

Poultry production has become the fastest growing livestock sector in the last three decades, with rapid intensification occurring in low- and middle-income countries (LMICs) and particularly in South and Southeast Asia [11]. In many of these countries, intensive production did not replace local farming and trading practices completely, resulting in multiple modes of production and distribution articulated in ways that are poorly understood and which vary according to market and other conditions. While such transformative changes have proven instrumental towards improving food security, nutrition and economic and societal development e.g. in China, India, Bangladesh among others, they also require careful monitoring and investigation. Indeed, the growth of poultry production and distribution networks has brought novel challenges in terms of disease management: intensive farming, limited surveillance infrastructure and veterinary services and in many examples poor biosecurity conditions [12,13] can lead to an environment replete with health hazards. For example, widespread sub-optimal use of antimicrobial drugs by poultry farmers represents a leading driver of the emergence of antimicrobial resistance [14–16].

In many LMICs, people prefer to obtain their poultry from live bird markets (LBMs), which are a longstanding feature of poultry trade and of urban and rural life. Within poultry PDNs, LBMs may be considered as hubs, sites wherein large numbers of people, and critically birds, meet and mix [17, 18]. Thus, they are major hotspots of AIV amplification and evolution [19], and have been implicated in sustaining viral transmission in domestic poultry [20]. The diverse ecology of AIV strains circulating within LBMs in Asia has been documented extensively [21–24]. Low pathogenic strains such as H9N2 AIV are commonly found among LBMs in Bangladesh, often at higher rates than in surrounding farms [25–27]. Since its first identification in 1996, highly-pathogenic H5N1 influenza has been detected in LBMs in many Asian countries [28–32].

While the biological risks within poultry production systems are widely acknowledged, they remain poorly characterised. This is partly due to the inherent complexity of PDNs, which makes it difficult to understand how such risks are modulated and increased along poultry value chains. Previous modelling efforts have focused on disease transmission within specific PDN settings, e.g. single farms or LBMs [33, 34], or some PDN segment, such as networks of farms or LBMs [18, 35–37]. Attempts to account for poultry or livestock PDN structure in infectious disease modelling are rare and mostly theoretical, often leaving out many epidemiologically relevant details of poultry production and distribution [38, 39]. Recent PDN mapping efforts have provided a clearer picture of PDNs in several Asian countries [40,41]. A central observation is that PDNs are highly heterogeneous across countries, poultry types, and even within the same country. Therefore, a better understanding can be achieved by extending and developing modelling to increase our understanding structural heterogeneities within and across PDNs.

To address this gap, we introduce EPINEST, a novel agent-based model (ABM) that allows simulation of pathogen transmission on top of realistic, empirically derived assumptions about poultry movements. EPINEST generates synthetic PDNs consisting of the key nodes, e.g. farms, traders, LBMs, that are responsible for the production and transportation of chickens through the PDN until they are sold to end-point consumers. Extensive data about farming and trading practices, collected mainly from field surveys, is used to inform PDN generation and simulation [18, 27]. Farm-specific data, for example, include farm locations, capacity and statistics of distinct stages of production cycles. Trader-level data encompass details of purchases and sales involving individual actors, origins of purchased poultry, and trader movements. EPINEST allows for substantial flexibility for users in terms of specifying PDN structure and functioning, making it a suitable framework to carry both data-driven and more open-ended analyses. In fact, the ABM permits customisation of many PDN properties, thus allowing users to explore a wide range of hypothetical PDN configurations.

This ABM provides a unified and flexible modelling framework to simulate epidemic dynamics in poultry PDNs and is the outcome of a wider interdisciplinary research initiative [42]. Within this context, EPINEST will enable investigating the amplification and dissemination of a wide range of health hazards, including AIV, *Campylobacter* and anti-microbial resistance genes in poultry systems in Bangladesh, India, Vietnam and Sri Lanka. More broadly, our framework may also be tailored to distinct poultry and livestock production realities to tackler a wider range of epidemiological questions.

In this paper, we provide a detailed description of our ABM and illustrate how to use it to explore a range of PDN structures and to better understand aspects of pathogen transmission in PDNs. The examples presented here are based on a broiler (chickens reared for meat) PDN in Bangladesh, which has been characterised extensively [18, 40], while epidemic simulations focus on the paradigmatic case of AIV transmission. The latter also illustrate an important feature of our framework, namely the ability to simulate multiple co-circulating pathogens and their interactions.

## Results

### Synthetic poultry networks

To address questions about the eco-epidemiological dynamics of AIVs and other poultry-related pathogens, we implemented an agent-based model to simulate pathogen transmission on top of synthetic PDNs. Within our framework, generated PDNs consists of four main types of nodes: farms, middlemen, vendors and LBMs (Fig. 1A). The system works as a supply chain where chickens are reared in farms starting from day-old chicks and are later transported to LBMs by middlemen (more details can be found in the Materials and Methods section and in Text S1). Once they arrive at the LBM stage, chickens are handled by vendors. These vendors may then sell chickens to other vendors operating in the same or different LBMs, and/or to endpoint consumers, in which case chickens are removed from the PDN. At any stage where chickens are exchanged, other than to the endpoint customer, an opportunity arises for pathogen exchange and mixing.

**Fig 1:**
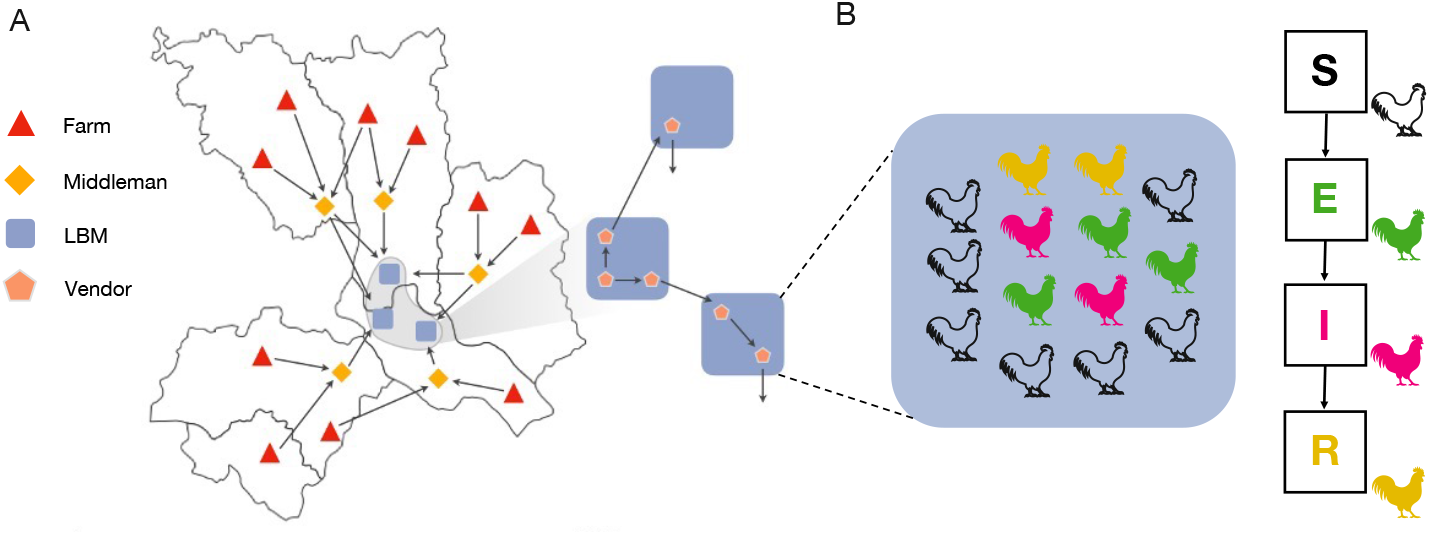
Model schematics. (A) Synthetic PDN and poultry movements. Chickens are produced in farms (red) across the study area, and transported to LBMs (blue) by middlemen (yellow). These are mobile traders that may collect chickens from multiple farms located in one or more upazilas/sub-districts (an administrative area below that of a district in Bangladesh). Within LBMs, chickens are handled by vendors (orange) and may be moved between LBMs as a result of vendors’ trading practices. (B) Individual settings associated with farms, middlemen, LBMs (when open) and vendors (overnight, when LBMs are closed) provide the context for pathogen transmission, under the assumption that chickens mix homogeneously within the same setting. The panel zooms in on a single LBM, where chickens are colour-coded according to disease status: susceptible (S), exposed or latent (E), infectious (I) and recovered or immune (R).

To illustrate the ability of the model to synthesize realistic poultry movements, we simulate a small PDN consisting of 1200 farms scattered across the 50 upazilas (sub-districts) that supply the largest amount of broiler chickens to LBMs located in Dhaka (Fig. 2A). The simulated PDN includes 20 distinct LBMs, 163 middlemen and 444 vendors, and allows the trade of chickens between LBMs. Numbers of middlemen and vendors can not be specified a priori; instead, they are determined dynamically by initially calculating the average number of chickens that are sold by farms to each LBM daily. These calculations depend on the spatial arrangement of farms, their sizes and frequency of selling, i.e. parameters that can be specified a priori. The capacity of each trader (middleman or vendor), i.e. the maximum amount of chickens that he/she can purchase daily is also fixed over the course of a simulation.

**Fig 2:**
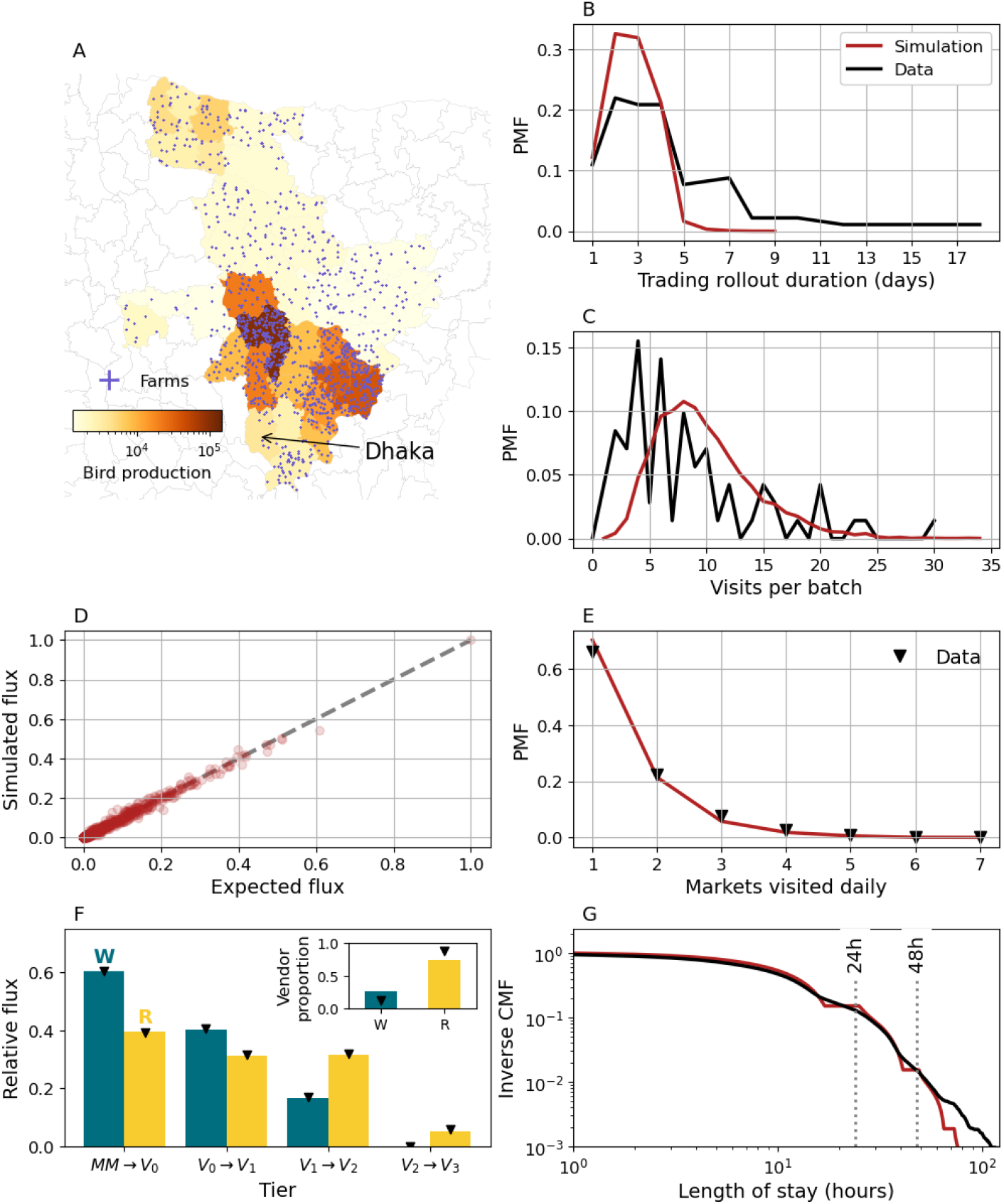
Simulating poultry movements. (A) Spatial population of 1200 farms supplying Dhaka. Farm locations are generated as described in Text S1 and assigned preferentially to upazilas with a larger observed outgoing chicken flux (colour scale). (B) Expected (black) and measured (red) distribution of times required to sell an entire batch. (C) Expected and measured distributions of transactions a single batch is split into. (D) Measured vs expected relative flux between individual pairs (dots) of upazilas and LBMs. (E) Distribution of LBMs serviced daily by individual middlemen. (F) Proportion of chickens sold to wholesalers (W, teal) and retailers (R, yellow) by LBM tier in simulations (bars) and data (markers). *MM* → *V*_0_ refers to transactions involving middlemen and first tier vendors, while *V*_*L*_ *→V*_*L*+1_ represents inter-tier transactions. For each tier, bars do not add up to 1 since wholesalers can sell to end-point consumers as well. Inset shows proportions of wholesalers and retailers. (G) Marketing time distribution. Results are obtained from a single simulation with default settings. Farm data are obtained from [27]. Data about middlemen and vendor trading practices and marketing times are obtained from [18].

Farms sell all their chickens at the end of a production cycle. The trading phase may require multiple days to complete and the flock may be split into multiple transactions involving different middlemen. Fig. 2B,C show that both the distributions of farm trading times and numbers of transactions per production cycle obtained through simulations are consistent with field observations. Upstream transportation and distribution of poultry operated by middlemen represent an important driver of poultry mixing in LBMs [18]. In simulations, middlemen direct previously purchased chickens to LBMs depending on where these have been sourced from. In practice, a chicken bought in upazila *a* is sold in market *l* with probability *f*_*a,l*_, as estimated from field questionnaires Fig. 2D shows that the ABM generates poultry fluxes between individual upazilas and LBMs that are in excellent agreement with the corresponding expected values (i.e. *f*_*a,l*_). Moreover, the allocation algorithm ensures that individual middlemen deliver chickens to a desired number of LBMs, as specified by some statistical distribution. The agreement between empirical and simulated frequencies of unique LBMs visited daily is shown in Fig. 2E. At the market level, wholesaling activities and vendor movements between LBMs further contribute to poultry mixing. Once a chicken enters an LBM, it may be sold multiple times to secondary vendors before reaching end-point consumers [18, 43]. In order to better capture the inner organization of LBMs, the model structures vendors in tiers according to their position along transaction chains (Fig. 2F). Finally, we show the realised distribution of poultry marketing times alongside another estimate obtained using a different approach [18] (Fig. 2G). Further statistics about individual actors and poultry transactions can be found in Fig. S1, S2 and S3.

Selected aspects of generated PDNs can be easily manipulated within our framework, allowing flexibility in exploring PDN configurations. In Fig. 3, for example, we examine different distributions of LBMs serviced (*Pr*(*k*_*m*_)) by individual middlemen on a daily basis (Fig. 3A). As we increase the number of LBMs serviced per middleman, ⟨*k*_*m*_⟩ on average, middlemen trade with more vendors (Fig. 3B); consequently, individual transactions involve fewer birds since the total cargo is the same (Fig. 3C). Fig. 3B also suggests that the small discrepancy observed in Fig. 3A at larger ⟨*k*_*m*_⟩ is due to the limited amount of vendors (inset).

**Fig 3:**
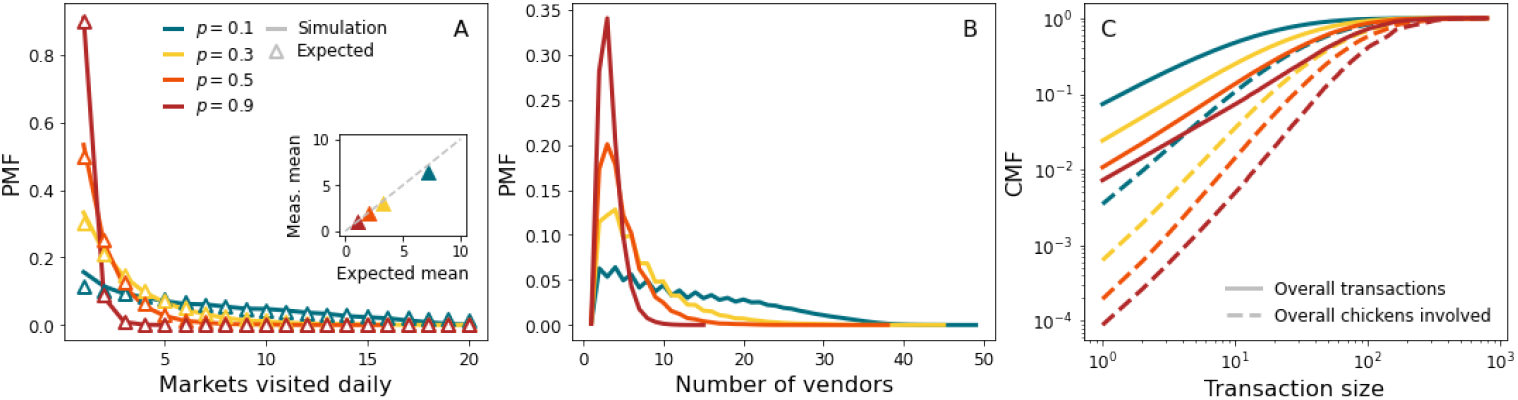
Markets serviced daily. (A) Expected (scatters) and measured (lines) distributions of markets serviced daily. Expected distributions are of the form 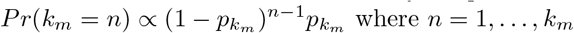. Inset compares expected and measured average numbers of markets serviced. (B) Distribution of vendors a single middleman trades daily with. (C) Cumulative distribution of sizes of transactions involving middlemen and vendors (solid lines). Dashed lines represent cumulative proportion of chickens sold in transactions up to a given size. Results are averaged over 50 simulations from 10 different PDN realisations.

We also present the impact of vendors’ trading practices on poultry marketing time. In particular, we alter the probability *p*_*empty*_ that a vendor sells its entire cargo in a single day, the fraction *ρ*_*unsold*_ of unsold birds in presence of some surplus (occurring with probability 1 −*p*_*empty*_). In addition, we consider high and low tendency to prioritise selling older (i.e. previously unsold) chickens over newly purchased ones. Varying parameters *ρ*_*unsold*_ and *p*_*empty*_ affects the average marketing time (Fig. 4A), as well as the proportion of chickens being offered for sale on multiple days (Fig. 4B). Full distributions of marketing times can be found in Fig. S4. Prioritizing the sale of older chickens had a negligible effect on these statistics. Indeed, prioritizing older chickens is compensated by a delay in selling newly purchased chickens (Fig. 4C).

**Fig 4:**
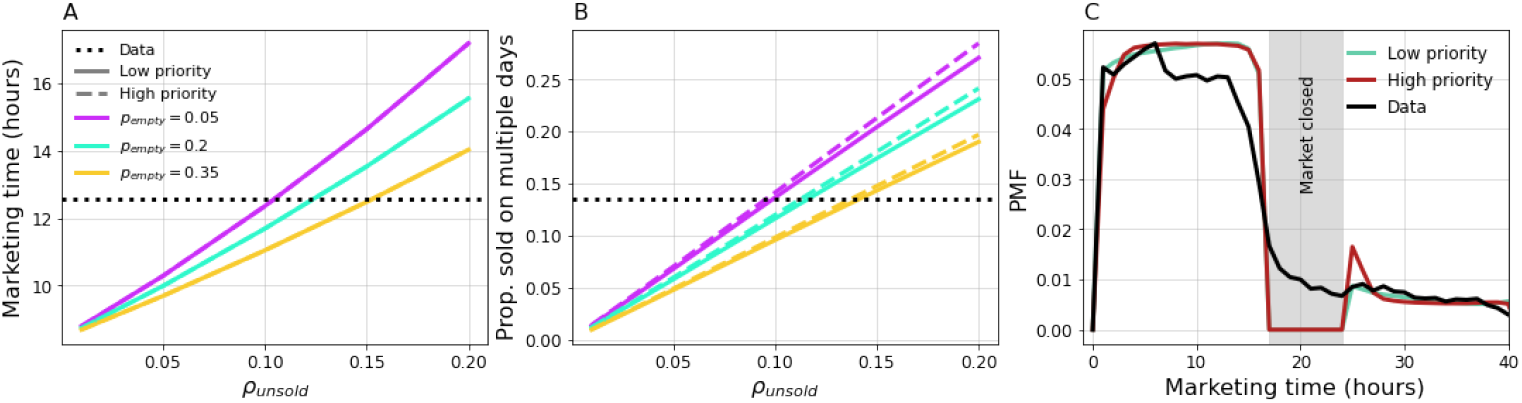
Vendor trading practices. (A) Average marketing time as a function of *ρ*_*unsold*_ for different values of *p*_*empty*_. Solid and dashed lines correspond respectively to low (10%) and high (90%) frequency of vendors prioritizing trading older chickens. (B) Proportion of marketed chickens offered for sale on multiple days. (C) Marketing time distributions for low and high frequency of vendors prioritizing older chickens. Here, *ρ*_*unsold*_ = 0.1 and *p*_*empty*_ = 0.2. Results are averaged over 50 simulations from 10 different PDN realisations.

As a final example, we examine the role of vendor movements between LBMs in promoting the mixing of chickens from different upazilas/sub-districts. Networks of LBMs defined by trader movements can vary considerably across poultry types, countries, and even cities within the same country [40]. In Chattogram, for example, vendors trading broiler chickens operate almost exclusively in a single market (Fig. 5A). In Dhaka however this is not the case, resulting in frequent vendor movements that are articulated in a top-down structure where central and peripheral markets can be identified (Fig. 5B). In fact, removing a single edge in the network shown in Fig. 5B is sufficient to make it acyclic, suggesting a hierarchical organisation.

**Fig 5:**
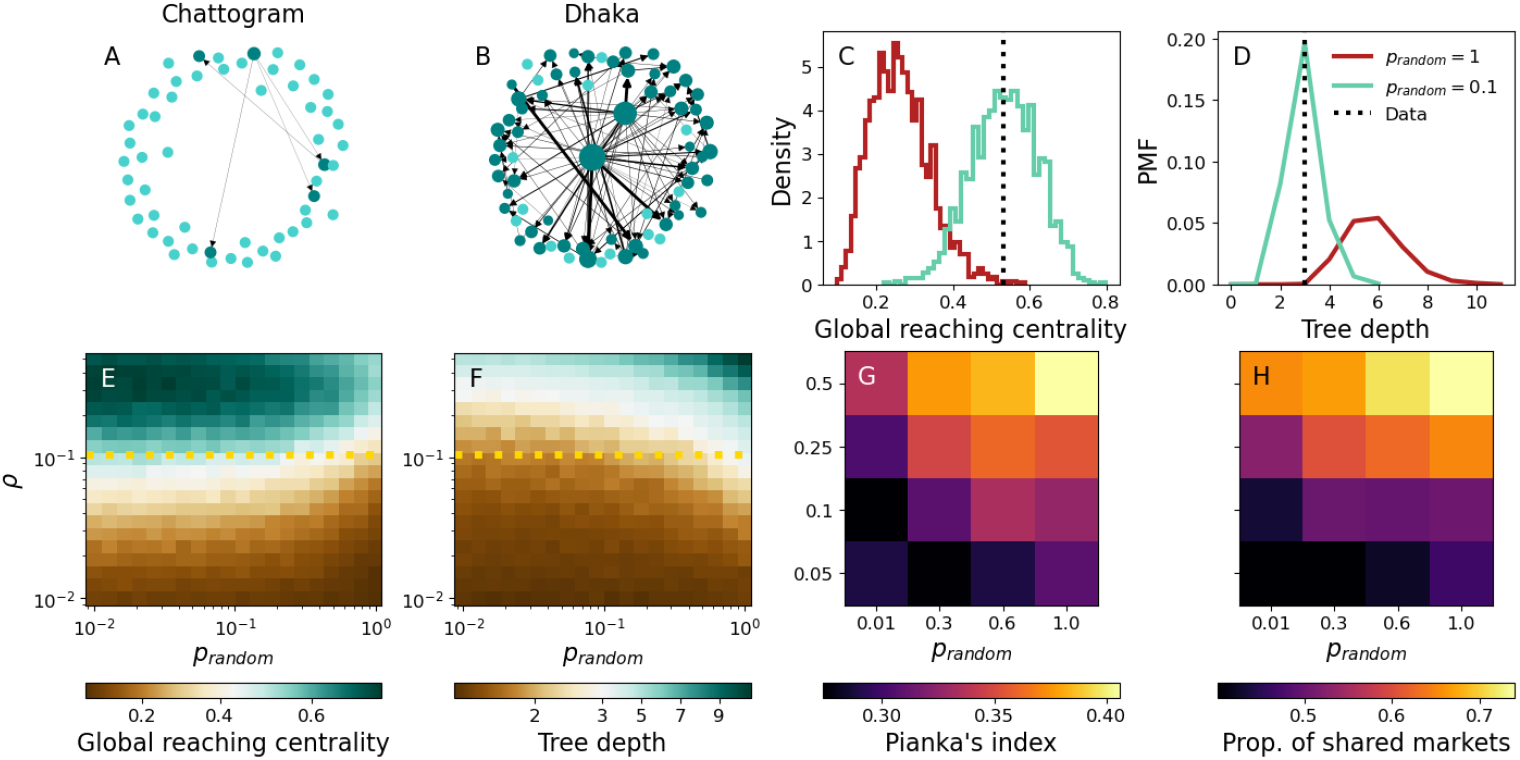
LBM networks and poultry mixing. (A,B) Broiler LBM networks for Chattogram and Dhaka, respectively. An arrow pointing from market *l* to *l*^*′*^ indicates at least one movement in that direction, while arrow thickness is proportional to the number of vendors moving on that edge. Node size is proportional to the outgoing weight, i.e. the total number of vendors leaving it. Isolated and connected nodes are shown in cyan and teal, respectively. (C,D) GRC and TD, respectively, for Dhaka’s network (line) and ensembles of 2000 synthetic LBM networks with the same density as Dhaka’s network and *p*_*random*_ = 1 (red) and *p*_*random*_ = 0.1 (cyan). (E,F) Average GRC and TD, respectively, across 100 networks with 20 nodes and as a function of *ρ* and *p*_*random*_. Dotted line denotes Dhaka’s density. (G,H) Pianka’s index of overlap and proportion of markets where it is possible to find chickens from different upazilas/sub-districts, respectively, as a function of network parameters. Performing the same measurement before any vendor movement occurs, yields an overlap (Pianka’s) of 0.261, and 25,7% shared markets, on average. This represents the baseline overlap due to middlemen sourcing chickens from farms and selling them to vendors. Results are averaged over 50 simulations from 10 different PDN realisations. All other PDN parameters are set to default values.

Within our framework, we encode inter-market mobility in a graph *G*, whose entries *G*_*i,j*_ represent the probability that a vendor purchasing in market *i* moves to market *j* (or remains in *i*) to sell. As outlined above, vendors are further arranged in tiers, so that vendors in tier *L* (*V*_*L*_) can only buy poultry from wholesalers located in tier *L−* 1 or, in the case of *L* = 0 vendors (*V*_0_) from middlemen trading in LBM *i*. For each vendor, purchase and sell locations remain fixed throughout a simulation.

To explore inter-market mobility, we use a generative network model to create mobility networks *G* akin to that of Fig. 5B. In practice, we generate directed acyclic graphs (DAGs) of varying density and amount of hierarchy (see Materials and Methods section), according to parameters *ρ* and *p*_*random*_. *ρ* represents the density of connections, while *p*_*random*_ is the probability of an individual connection emanating from a source LBM that is selected randomly, rather than proportionally to their actual number of connections. To quantify the degree of hierarchy in a DAG *G*, we measure its global reaching centrality (GRC, see Fig. 5C,E) and tree depth (TD, see Fig. 5D,F). GRC measures how well every node can reach other nodes in the network with respect to the most influential node; it takes value 1 in the case of a star graph and approaches 0 when all nodes have the similar influence (no hierarchy). In contrast, TD represents the longest directed path in *G*. Hierarchical DAGs, e.g. stars, tend to be more compact and hence shallower than random structures. Setting *p*_*random*_ = 1 yields DAGs with little hierarchy, as edges are allocated randomly. In contrast, *p*_*random*_ → 0 introduces additional structure. Fig. 5C,D show GRC and TD, respectively, for Dhaka’s network and for DAGs generated with *p*_*random*_ = 1 (red) and *p*_*random*_ = 0.1 (cyan) while keeping the density of edges constant. Clearly, Dhaka’s network is significantly more hierarchical and compact than random DAGs; in contrast, DAGs generated with *p*_*random*_ = 0.1 provide a much closer fit in terms of both GRC and TD.

We also summarize framework output by quantifying poultry mixing across 20 LBMs for different combinations of *ρ* and *p*_*random*_ (GRC and TD are shown in Fig. 5E,F, respectively). Here mixing refers to the extent to which chickens from distinct regions are brought together within LBMs. Upstream distribution, managed by middleman, and vendor movements between LBMs are the factors driving chicken mixing within this model. To quantify the amount of mixing, we record the geographic origins of chickens offered for sale in each LBM and use Pianka’s index [44] to make pairwise comparisons of poultry populations marketed in distinct LBMs. Mean Pianka’s index values are shown in Fig. 5G as a function of parameters *ρ* and *p*_*random*_. Values close to 0 imply low overlap, while a value of 1 corresponds to identical distributions of geographic sources of poultry. Fig. 5H shows another, complementary quantification of poultry mixing in terms of the mean number of LBMs where it is possible to find chickens from two randomly chosen upazilas/sub-districts. In general, we find that chicken mixing increases with network density, while hierarchy has the opposite effect: random vendor movements are more effective at mixing chickens within this simplified network model. It should be noted that high levels of mixing can be observed even in the absence of vendor movements due to upstream distribution (overlap between the catchment areas of LBMs, of which middlemen are responsible; see caption of Fig. 5 for further details).

### Epidemic dynamics

In this section, we illustrate how our framework can be used to simulate and characterise pathogen transmission across PDNs. We first consider a single, AIV-like pathogen whose dynamics is described by a Susceptible-Exposed-Infectiou Recovered (SEIR) model, as depicted in Fig. 1B: upon infection, susceptible (S) chickens enter an intermediate exposed stage (E) and become infectious (I) after a short latent period *T*_*E*_ = 6 hours. Infectious chickens recover (R) after an infectious period *T*_*I*_ = 48 hours and become immune to further infection. Importantly, we assume that chickens do not die due to the disease. We assume that the pathogen (repeatedly) emerges at rate *α* in farms due to external factors (e.g. contacts with wild birds) and spreads through the PDN through a combination of poultry movements, intra- and inter-farm transmission (see Material and Methods section).

Model output comes at different levels of aggregation. Fig. 6A shows for example daily incidence within LBMs during the first stages of an outbreak. At the most granular level, individual transmission events and their metadata can be tracked as well. Using this information, we can reconstruct transmission chains originating from individual introduction events and characterise their spatiotemporal evolution (Fig. 6B). Fig. 6C further characterises farm outbreaks by summarising attack rates by production cycle.

**Fig 6:**
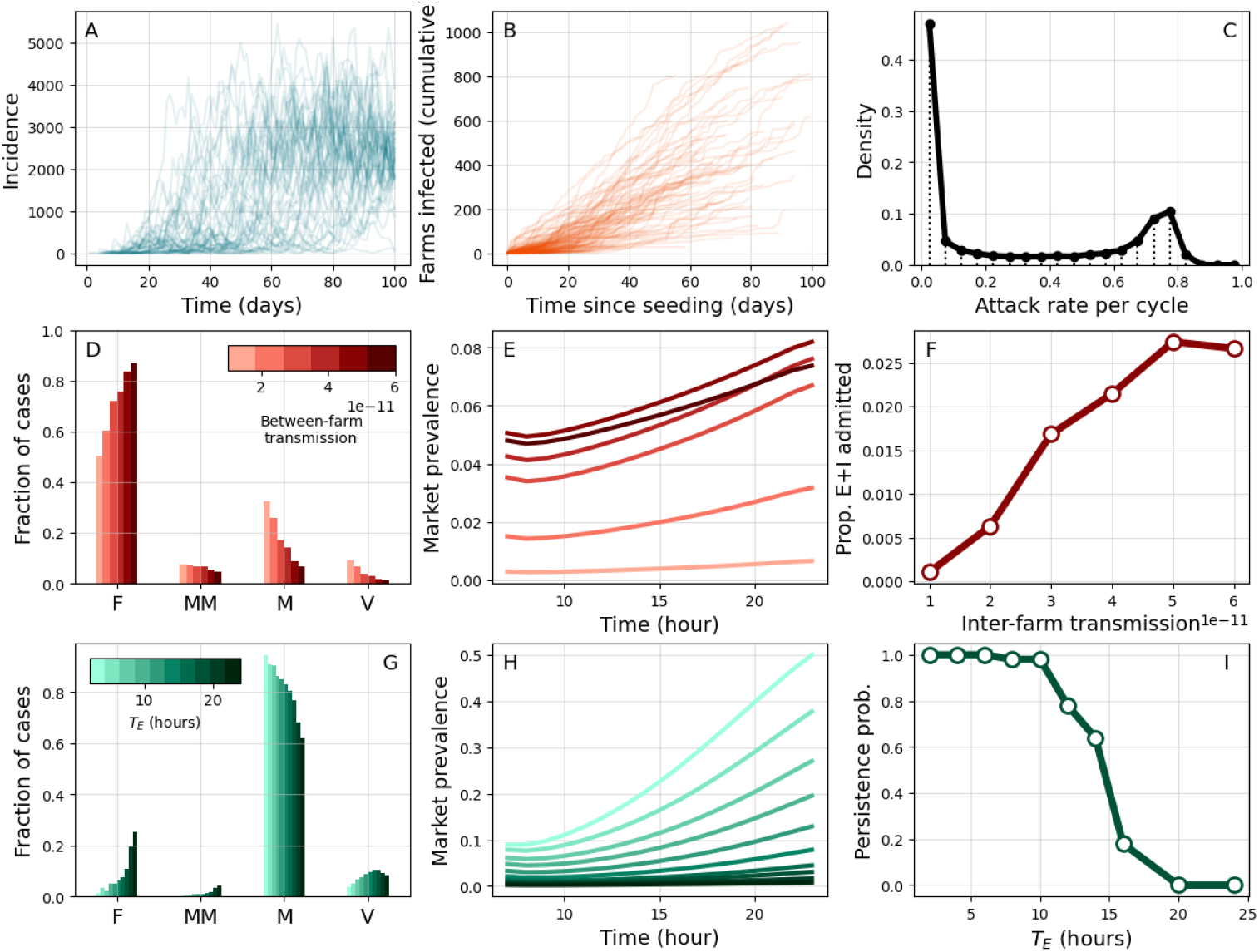
Epidemic dynamics. (A) Daily incidence in LBMs in multiple simulations. (B) Cumulative number of new farms infected over time from multiple clusters. Each cluster is initiated by a different infectious seed. (C) Distribution of attack rates for individual production cycles, conditional on at least one infection. (D-F) High farm transmission scenario (*w*_*F*_ = 0.2, *w*_*M*_ = 0.7). Colour scale corresponds to varying levels of inter-farm transmission *β*_*F F*_. (D) Proportion of incident cases in different setting types (F: farms, MM: middlemen, M: markets, V: vendors). (E) Average hourly prevalence in LBMs at stationariety. (F) Proportion of latent and infectious chickens entering markets daily as a function of *β*_*F F*_. (G-I) High LBM transmission scenario (*w*_*F*_ = 0.1, *w*_*M*_ = 2.4). Colour scale corresponds to varying latent period *T*_*E*_. (G,H) mirror (D,E). (I) Persistence is measured as the proportion of simulations where at least one transmission chain persisting in markets and vendors for longer than 50 days was observed. Results are qualitatively the same under different different criteria about the duration of transmission chains (Fig. S5). Other parameters are set to default values. Results are based on 50 simulations from 10 different synthetic PDNs.

In Fig. 6D-F, we investigate the role of spatial transmission in an endemic context. We do so in a scenario where most transmission events occur within farms (Fig. 6D), while viral amplification in LBMs is limited (Fig. 6E). Here, spatial transmission is a crucial factor in determining global levels of infection. Increasing the strength of inter-farm transmission *β*_*F F*_ facilitates spatial invasions, thus leading to more outbreaks on farms and infections (Fig. 6D). This results in an increasing number of infected chickens pouring into LBMs from farms (Fig. 6F), explaining also the increase in within-market prevalence observed in Fig. 6E.

Another important epidemiological question is whether AIV is transmitted and maintained in LBMs despite short marketing times. We address this question by considering an alternative endemic scenario where transmission is contributed mostly by LBMs (Fig. 6G). We find that a major limiting factor to viral amplification in LBMs is represented by the latent period *T*_*E*_ (Fig. 6H): delaying the onset of infectiousness corresponds to a shorter window of opportunity for transmission under short marketing times. In order to further demonstrate this point, we quantify persistence of transmission chains within LBMs (Fig. 6I). As *T*_*E*_ increases, opportunities for transmission are diminished and chains of infection stutter, leading to reduced persistence. In this case, the presence of AIV in LBMs can only be maintained through repeated introductions of infected poultry.

### Simulating multi-strain pathogens

Genomic surveillance in LBMs routinely identifies AIV lineages with distinct genetic signatures [45]. In some instances, the presence of multiple AIV subtypes, including the highly pathogenic H5N1 AIV, is also reported. Understanding this diversity requires, however, accounting for multiple, potentially interacting strains/pathogens that co-circulate in the same PDN. In this section, we use our framework to perform multi-strain simulations in a variety of PDN structures.

We illustrate this in Fig. 7, which shows SEIR simulations with 50 cocirculating strains. For simplicity, we assume that these share the same epidemiological parameters, namely *T*_*E*_, *T*_*I*_ and *β*, and generate partial cross-immunity after a single infection.

**Fig 7:**
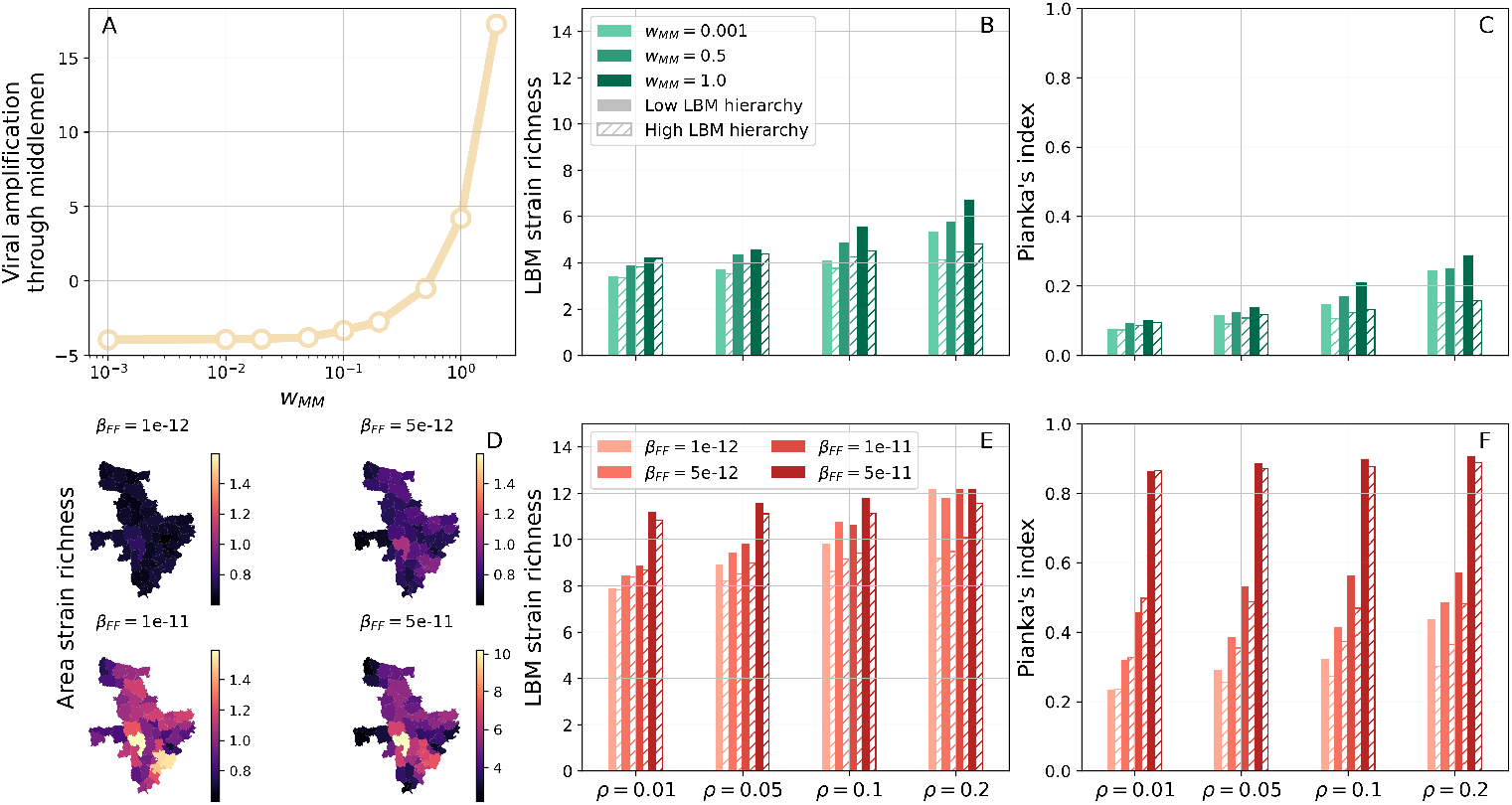
Multi-strain dynamics and viral mixing in LBMs. (A-C) Simulations with no inter-farm transmission (*β*_*F F*_ = 0). (A) Viral amplification happening through transportation from farm to LBM gates as a function of middlemen-specific transmission weight *w*_*MM*_. This is quantified through the difference between total numbers of exposed and infected chickens sold to vendors and purchased daily by middlemen. (B) Average strain richness (i.e. number of strains) in single LBMs as a function of density *ρ* of vendor movements (on the x-axis), *w*_*MM*_ (from light to dark). Solid and striped bars correspond to low and high hierarchy in vendor movements, respectively. (C) Average Pianka’s index of overlap between pairs of LBMs in terms of their catchment areas. (D-G) Simulations with inter-farm transmission. (D) Average richness per upazila for increasing *β*_*F F*_. Note that the bottom-right map uses a different colour scale. (E,F) Same as (B,C) but for varying *β*_*F F*_ and with *w*_*MM*_ = 0.001. We set *w*_*F*_ = 0.1 in (A-C) and *w*_*F*_ = 0.2 in (D-F), while *w*_*M*_ = 2.4 and *w*_*V*_ = 1 in all panels. Cross-immunity reduces susceptibility to secondary infections to *σ* = 0.3. Results are averaged over 50 simulations from 10 different synthetic PDNs. In each simulation, statistics are collected for 100 days after an initial transient of 2000 days.

Our aim is to measure the extent to which PDNs mix viral lineages from distinct geographical regions. To this end, we modify the external seeding protocol so that strain *s*_*i*_, *i* = 1, …, 50 can emerge only from upazila *i*. First, we investigate the role of viral amplification during the transport segment, which is operated by middlemen (Fig. 7A-C). To better disentangle the role of these actors, we consider low within-farm transmission and prevent inter-farm transmission fully by setting *β*_*F F*_ = 0. Consequently, viral mixing can not occur until chickens from different upazilas/sub-districts are collected by a middleman. As shown in Fig. 7A, increasing transmission during transport by varying *w*_*F F*_ leads to more infected chickens being introduced in LBMs, i.e. it results in viral amplification. Note, however, that below a certain value of *w*_*MM*_, middlemen may introduce fewer infections in LBMs than those they picked up at farms. Increasing *w*_*MM*_ has a modest positive effect on the average number of strains circulating (i.e. strain richness) in individual LBMs, and on the overlap between LBMs in terms of circulating strains (light to dark bars in Fig. 7B and C, respectively). We also explored, for fixed *w*_*MM*_, the role of inter-market mobility on these metrics. In this context, the density *ρ* of vendor movements had a positive effect on both strain richness and overlap between LBMs as it is promoting the dissemination of multiple strains across LBMs. In contrast, a larger degree of hierarchy in movements (striped bars) had the opposite effect, in agreement with findings from Fig. 5.

Finally, we consider a further scenario in which transmission within and between farms plays a central role in shaping epidemic dynamics, while transmission occurring during transport is assumed to be negligible (*w*_*MM*_ = 0.001). We find that increasing between-farm transmission *β*_*F F*_ leads to a wider spatial dissemination of strains even outside their upazila of origin (Fig. 7D). Consequently, a more diverse set of strains is supplied to LBMs, as evidenced by the number of strains observed at these locations (Fig. 7E). Also, because larger values of *β*_*F F*_ promote strain dispersal across the entire area, LBMs are now more similar to each other in terms of their strain populations (Fig. 7F). It should be noted, however, that increased within-farm transmission is responsible, at least in part, for the larger strain numbers and overlap between LBMs observed in Fig. 7E,F with respect to panels B,C. Finally, we note that the effects of density and hierarchy of vendor movements on ecological metrics are analogous to those observed in the previous scenario. These results are robust to increasing cross-immunity between strains (Fig. S6).

## Discussion

In this paper we have introduced EPINEST, an agent-based model to simulate the transmission of generic health hazards in the context of realistic poultry or livestock movements within a defined PDN. To the best of our knowledge, this work represents the first attempt to account for the structural complexities of poultry PDNs in the context of epidemic transmission modelling. Our model allows to generate synthetic PDNs consisting of key actors and settings involved in poultry production and distribution, namely farms, middlemen, LBMs and market vendors. Using Bangladesh as a case study, we illustrated the ability of our framework to reproduce empirical features of a broiler PDN. We used extensive data from field surveys to inform most aspects of the model, including farming and trading practices of key actors [18, 27, 46]. At the same time, our model offers the possibility to easily manipulate most properties of the network, allowing exploration of alternative PDN configurations. Importantly, we emphasize that our model may be applied to other contexts, e.g. different poultry types and countries for which sufficient data is available.

One of the main purposes of EPINEST was to assess the impact of PDN structure and stakeholders’ trading practices on pathogen transmission. For this reason, we prioritised including PDN components with the highest relevance to transmission dynamics. These include, for example, the time spent by chickens at different locations. All-in/all-out production, which is commonly implemented in commercial broiler farms, results in relatively homogeneous rearing times across farmed chickens, although these vary considerably among different chicken types. In contrast, LBMs are characterised by a much faster turnover, with most chickens being sold within a few hours and unsold chickens remaining for up to a few days. Longer marketing times are a well-established risk factor for AIV infection in LBMs, and have been linked to AIV persistence in these settings [18, 47]. To account for heterogeneity in marketing times, we explicitly account for a fraction of chickens being offered for sale on consecutive days.

Further basic ingredients of the model are the spatial distribution of poultry farms and their sizes. Both elements are highly relevant to disease transmission. Heterogeneities in farm locations can affect systemic vulnerability to epidemics and pathogen dispersal patterns [48–50], while higher livestock densities are associated with increased intra-farm transmission and may favor the emergence of virulent pathogens [9, 51]. In the absence of accurate data about farm locations, we generated random farm distributions complying with reported volumes of poultry production at the upazila level, and used field surveys to assign farm sizes [18, 27]. Nonetheless, we stress that our model can accommodate any distribution of farms. These may represent not only higher-resolution data, but also outcomes from more accurate generative models [52–54].

Our model also allows to control the degree of mixing of chickens along distribution and trading channels. The ability of PDNs to mix large numbers of chickens, particularly within LBMs, is well-established. The inter-mingling of different types of birds from potentially distant locations is concerning when associated to co-circulation of genetically distinct viruses. A recent phylodynamics study found substantial genetic structuring of H9N2 AIV by city in Bangladesh [55], compatibly with low overlap between the corresponding supplying production areas [18]. In contrast, viral lineages appeared to be highly mixed across LBMs within the same city, possibly indicating frequent connections between these markets. Live poultry trade has also been shown to be an important driver of regional AIV dissemination in China [56]. Here, poultry mixing within and between LBMs is dictated by two factors: first, upstream distribution via middlemen connects LBMs with farmed populations from a wide geographic area. Within our framework, geographic fluxes between regions (upazilas/sub-districts) and LBMs are expressed as a matrix that can be informed using field surveys or traceability systems. Second, wholesaling activities and vendor movements further contribute to stirring marketed poultry across LBMs. In this manuscript, we used a generative model to sample inter-market mobility networks, and quantified their impact on poultry mixing. We emphasize that more complex mobility patterns, informed either from data or through simulations, can be easily embedded within our framework.

A major feature of our model is that it allows simulating pathogen transmission while accounting for the complexity of poultry movements and PDN structures. Importantly, the epidemic layer is fully uncoupled from PDN generation. Thus, while current code supports simulations of SIR and SEIR dynamics only, implementing additional epidemic models is a relatively straightforward task. We illustrated how our ABM can be used to model AIV dynamics in both epidemic and endemic settings. In the former case, the ABM makes it possible to map the early dissemination of, e.g., an emerging AIV strain across farms and the rest of the PDN. The second scenario would be more suitable to describe endemic circulation of AIVs. In this context, relevant scientific questions that could be addressed using our framework include understanding how and where an endemic AIV is maintained and amplified along the PDN.

A novel aspect of our ABM is that it enables simulations of multiple cocirculating pathogens/strains and their interactions. This paves the way for a number of eco-epidemiological applications. As an example, we assessed the potential of PDNs to mix viral lineages originating from distinct geographical areas. Additional applications may consider the joint dynamics of endemic and emerging AIVs and simulate the early transmission dynamics of, say, highly-pathogenic H5N1 AIV against a background of (cross-)immunity generated by endemic circulation of H9N2 AIV [57].

As any modelling framework, there are limitations to our ABM. Despite our efforts to account for the structural complexity of PDNs, our focus on epidemiological investigations meant that several aspects of real PDNs could not be included in the model. For example, actors’ behaviours are treated as fixed parameters external to, rather than emerging from, the dynamic system being modelled. In reality, the decisions made by individual traders to sell or purchase birds is influenced by social, economic and epidemiological factors. These may include uncertainty about market conditions and fear spurred by disease outbreaks [58, 59]. In addition, unequal power dynamics often constrains trading ties [40, 60]. In this context, we plan on expanding our ABM’s capabilities to include simple reactive behaviours, e.g. farmers selling chickens pre-emptively following a surge in bird mortality [59]. Other extensions could include mixing of different poultry species, different farming systems and trading practices, such as second-line middlemen purchasing chickens from other traders, and different biosecurity measures implemented at different LBMs to limit pathogen spread. Finally, although we wrote our model in C++ to improve simulation speed, computational constraints make it difficult to scale up simulations to more than a few millions of farmed chickens. This is a common challenge in agent-based models, where the increased amount of detail is traded off by computational costs.

In conclusion, we implemented a novel agent-based model to jointly simulate realistic poultry movements and epidemic trajectories. Realised structures encompass a wide-range of PDN configurations as encountered in many countries in South and Southeast Asia, and potentially even other livestock production systems with similar structure to the one discussed here. Compared to existing ABMs devoted to veterinary epidemiology applications [61–64], ours offers the ability to run both single- and multi-strain simulations. In addition, the simulator can be programmed to yield a wide range of outputs, including individual transactions and chains of infections, hence providing a full characterisation of the underlying system. This model is a unique tool in the One Health context as it allows investigation of a range of epidemiological scenarios and helps us to understand better the role of different structural aspects on disease transmission. Immediate applications of this model will allow exploration of the transmission and amplification of AIVs and anti-microbial resistance genes within poultry PDNs.

## Materials and Methods

### Generating synthetic PDNs

In general, a PDN denotes the ensemble of actors that are involved in the production and/or distribution of a product such as poultry and their interactions. At any point in time, a chicken is physically located within one and only one setting, such as a farm, a middleman’s truck, an LBM or a vendor-owned shed during the night.

Our generative algorithm instantiates a population of actors based on external specifications. First, a spatial distribution of farms must be provided alongside the corresponding geographic setup. The latter consists of a partitioning of the study area into a set of non-overlapping regions. In this study, we take upazilas/sub-districts as regional units. Second, the user specifies a number of LBMs and their catchment areas. In practice, this is achieved by specifying a matrix *f*_*a,l*_ representing the relative fluxes of chickens reaching market *l* = 1, …, *N*_*M*_ from area *a* = 1, …, *N*_*A*_. A full description of LBMs requires a set of weights *w*_*l,l*_^*′*^ encoding the probability that a vendor purchases chickens in LBM *l* and trades in LBM *l*^*′*^ (with possibly *l* = *l*^*′*^). Finally, a number of parameters influencing farming, distribution and trading practices should be specified as well (these are described in Text S1). With these details, the algorithm computes the expected poultry fluxes between farms and LBMs and allocates enough vendors and middlemen to satisfy such demand. At the LBM stage, vendors are allocated in a tier-wise fashion depending on the volume of chickens supplied by middlemen, inter-market movements, and wholesaling practices. Eventually, it is possible to generate more middlemen and vendors than strictly required based on heuristic calculations by inflating the expected supply of chickens handled by middlemen and vendors through multiplicative factors *ϵ*_*MM*_ and *ϵ*_*V*_.

### Modelling inter-market movements

As detailed in Text S1, a vendor purchasing chickens in LBM *i* is assigned to trade in LBM *j* with probability *G*_*i,j*_.

We sample the weights *G*_*i,j*_ from a generative network model defined by a growth mechanism: we add LBMs *j* = 1, …, *N*_*M*_ one at a time and establish links *i* → *j, i < j* as follows: first, we draw the number of incoming edges (in-degree) *z*_*j*_ ∼ *Binomial*(*ρ, j* − 1). Second, we sample *z*_*j*_ LBMs (with *i < j*) without replacement either at random with probability *p*_*random*_, or proportionally to their out-degree. This yields a binary matrix ℐ_*i→j*_ denoting existing connections. Weights are then calculated as:

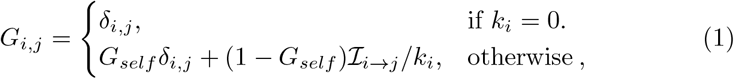

where *G*_*self*_ is the probability of a vendor operating in a single market *i*, conditional on the out-degree *k*_*i*_ = Σ_*j*_ ℐ_*i→j*_ being positive. Here we set *G*_*self*_ = 0.8.

### Global reaching centrality

GRC is defined based on the notion of local reaching centrality *C*_*R*_(*i*), which quantifies the proportion of nodes reachable from node *i* through directed edges. Based on this definition, we calculate GRC by subtracting *C*_*R*_(*i*) from the maximum observed value 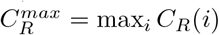, and averaging over all nodes:

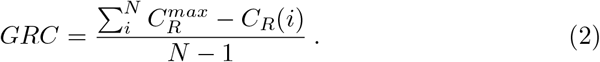

### PDN dynamics

Within simulations, actors follow a daily routine. Let *t* = 0, …, 23 indicate the time of the day (each time step is 1 hour long). Unless otherwise stated, default parameters indicated in Text S1 are considered. LBMs open between *T*_*open*_ and *T*_*close*_; at *t* = *T*_*open*_, vendors move to LBMs, followed by middlemen. Middlemen then proceed to sell their cargo to frontline vendors, i.e. those in the first LBM tier (*L* = 0). In the next time step (*t* = *T*_*open*_ + 1), some of these move to another LBM and trade with second-tier vendors, who in turn sell chickens to vendors in the tier after that, repeating the process until the last tier is reached. Vendor movements and wholesaling are therefore resolved sequentially, in a tier-wise fashion, at time *t* = *T*_*open*_ + 1. In contrast, retailing activities roll out between *T*_*open*_ + 1 and *T*_*close*_. At *T*_*close*_, both wholesalers and retailers leave LBMs alongside any unsold chickens. Overnight, these chickens are stored in some other place, e.g. in a shed. Importantly, all chickens from the same vendor are stored in the same place.

At some time *t* = *T*_*farm*_ we update farms: empty farms may recruit a new batch of chickens, while active farms may offer birds for sale depending on batch age. After that, always at *t* = *T*_*farm*_, middlemen are updated: first, they decide whether to cover a different set of upazilas/sub-districts. Then, they contact farms within covered upazilas/sub-districts in order to purchase chickens. At this stage, middlemen only determine how many chickens to collect from each farm; the collection may happen anytime between *T*_*farm*_ and *T*_*open*_ on the following day.

### Epidemic dynamics

In this work, we consider a general transmission model involving a generic number of strains. Each strain, indexed by *s*, spreads according to SEIR dynamics. Infected chickens become infectious only after a random latent period 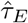, sampled from a distribution 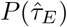 with mean *T*_*E*_. Analogously, infectious chickens recover after a random time 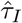, sampled from a distribution 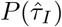 with mean *T*_*I*_. All epidemiological parameters are listed in Text S1.

Here, transmission is assumed to occur through infectious contacts among chickens from the same setting. Other transmission mechanisms, including external introductions and inter-farm transmission, are described in Text S1. During a time step, an infectious chicken *i* contacts a single chicken *j*, chosen at random within the same setting, and transmits strain *s* with probability:

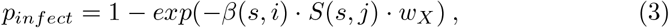

where *β*(*s, i*) transmissibility of strain *s* and *S*(*x, j*) is susceptibility of chicken *j* to *s*. The factor *w*_*X*_ is a multiplier that depends only on the underlying setting type (F,MM,M,V).

In general, the transmission rate *β*(*s, i*) and susceptibility *S*(*s, j*) may depend on the immune state of infector and infectee, respectively. Importantly, different functional forms of *β*(*s, i*) and *S*(*s, j*) embody different assumptions about immune cross-reactions induced by previous exposure to other pathogens/strains. In this work we consider uniform transmission *β*(*s, i*) = *β*, irrespective of immune state, and susceptibility *S*(*s, j*) = 0, 1, *σ* depending on whether *j* has already been infected with *s*, is fully naive or was infected with some other strain *s*^*′*^ ≠ *s*, respectively. The parameter *σ* ∈ [0, 1] represents reduced susceptibility due to cross-immunity, and interpolates between sterilising cross-immunity (*σ* = 0) and no cross-immunity (*σ* = 1).

## Supporting information

### Data availability statement

Data and code to generate and analyse simulation output are available at https://github.com/francescopinotti92/EPINEST.

## Supplementary Text for

### 1 Data analysis

#### 1.1 Farmers

##### Questionnaire data

Data were generated through a cross-sectional study that collected information about farming practices from 100 distinct farms in Chattogram division,

Bangladesh [1]. Available data includes farm sizes, the number of production cycles completed in one year, as well as the numbers of traders and transactions involved in clearing individual batches (Fig. A1). Here we focus on the 47 farms that reported trading broilers and raising a single batch at a time in a single shed. It should be noted, however, that while the questionnaire aimed to uncover farmers’ practices over one year, it is not possible to know whether farmers raised at most one batch at a time for the entire period as their answers reflect their situation during the interview.

##### Farm/batch size

We modelled batch sizes *S*_*F*_ by assuming a truncated negative binomial distribution. More in detail, we assumed:

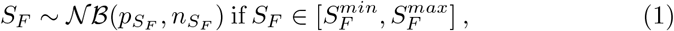

where 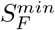, 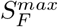 are the minimum and maximum observed batch sizes. For simplicity, we estimated parameters 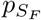, 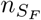 by maximum likelihood without accounting for truncation and verified a posteriori that the resulting bias was small. Note that for this type of farms, batch size is the same as farm size, since they do not rear multiple batches at the same time.

##### Replenishment time

We modelled replenishment time *τ*_*replenish*_ by assuming a shifted negative binomial distribution, i.e.:

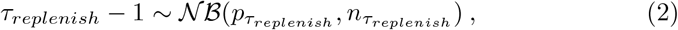

#### 1.2 Middlemen

##### Individual middlemen properties

We used data from questionnaires to inform middlemen’ trading patterns. Consistently with the main manuscript, we restrict our analysis to broiler data only [2].

##### Number of chickens bought daily

We divided the total number of chickens bought during the entire study period by each middlemen with the number of days in which he/she purchased any poultry. We used these raw values to construct a discrete distribution from which to sample middlemen batch sizes *S*_*MM*_ in the ABM.

##### Number of markets visited daily

As a first step, we evaluated the number of working days for each middlemen by combining sale frequencies of different breeds. This was straightforward for middlemen that sold chickens every day during the study or sold a single chicken breed. One middleman declared selling chickens 4 times in a week; in this case, we set the number of working days to 4. We then obtained the mean number of daily market visits *k*^(*daily*)^ by dividing the total number of market visits made by each middleman during the study period with the corresponding number of working days. Note that some values *k*^*daily*^ are not integer numbers due to the previous calculation. Finally, we used maximum likelihood to fit a geometric distribution with parameter 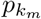 to data 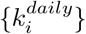, with *i* ranging from 1 to the total number of middlemen interviewed. More precisely, the maximum likelihood estimate for 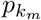 is given by the inverse of the sample mean of data 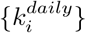. This formula works also when some values 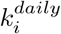 are not integer numbers (Fig. A2).

**Figure A 1:**
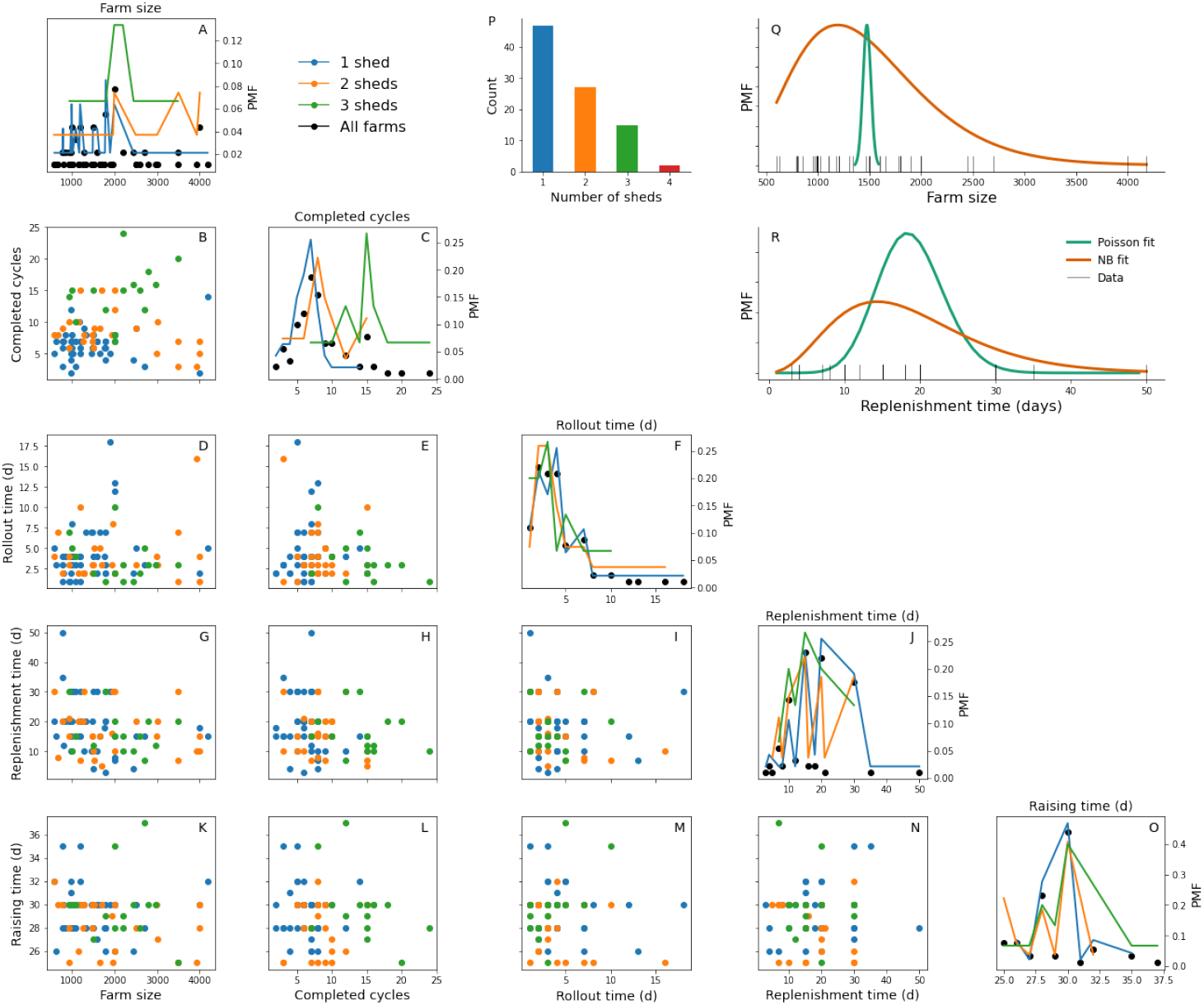
Farm statistics from survey data and statistical fits. (A-O) Univariate distributions (diagonal) and pairwise scatter plots (off-diagonal) for several farm statistics, coloured by the number of available sheds. Shown quantities include farm size (A), completed production cycles per year (C), trading rollout duration (F), time to replenish the farm (J) and raising time (O). We did not show farms with 4 sheds, as there were only two (see panel P showing frequencies of sheds per farm). (Q) Poisson (green) and truncated negative binomial (orange) fits to batch size data (black ticks). (R) Poisson (green) and shifted negative binomial (orange) fits to replenishment time data (black ticks). In both cases, a negative binomial distribution provides a better fit than Poisson to underlying data. Analyses in (Q,R) were restricted to farms with a single shed.

**Figure A 2:**
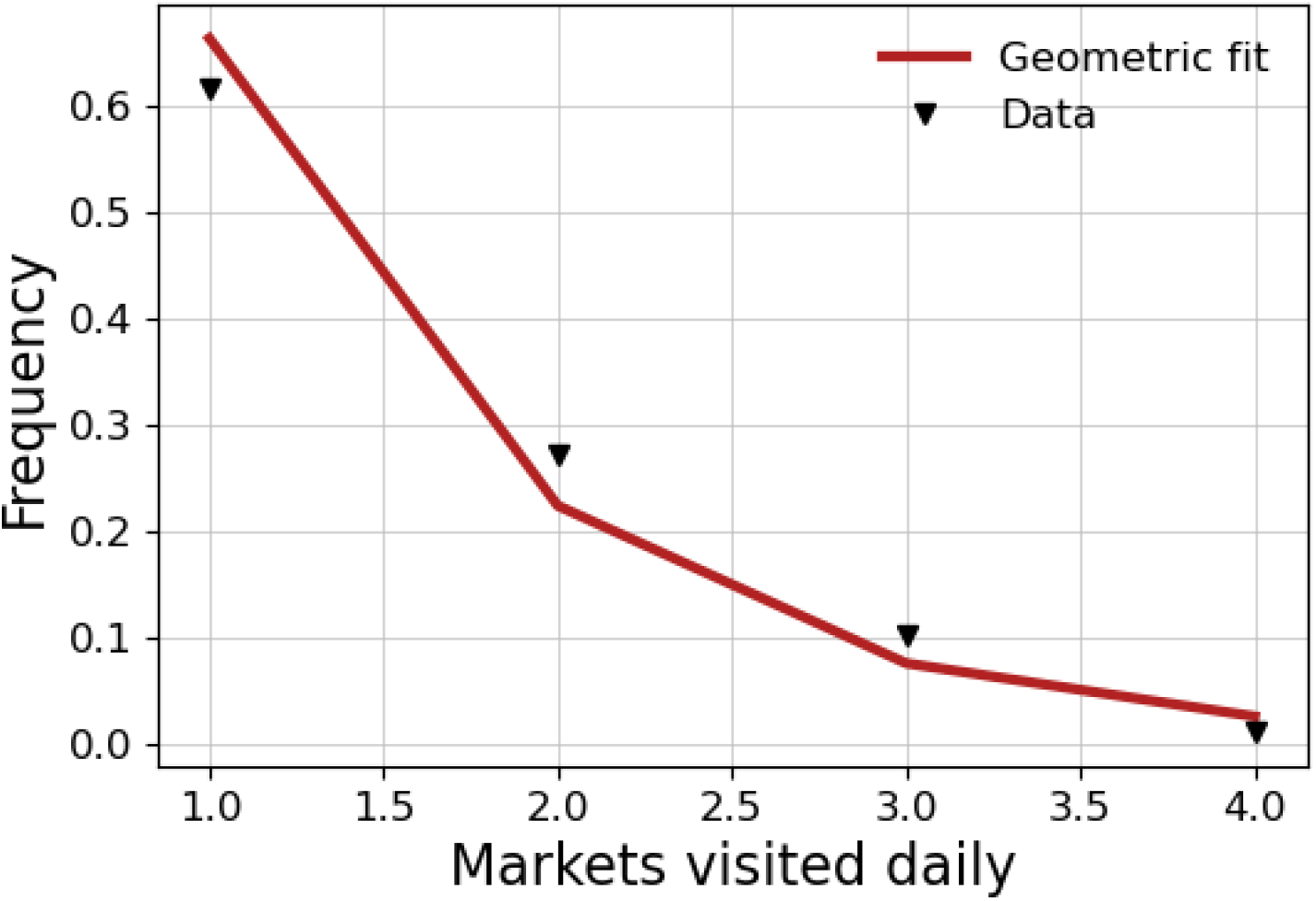
Distribution of daily market visits. Fitted (line) and observed (markers) frequencies of markets visited daily. The fitted distribution is geometric with parameter *p* estimated using maximum likelihood. Note that some data points were not integers and were therefore rounded to the nearest integer in this plot. The fitted distribution corresponds to black markers in Fig. 2E, but is here shown up to value 4, i.e. the maximum observed count.

#### 1.3 Vendors

##### Individual vendor properties

We used data from questionnaires to inform vendors’ trading patterns. Unless otherwise stated, we repeat every analysis for retailers and wholesalers.

##### Number of chickens bought daily

We first reconstructed the number of chickens bought by a single vendor over the survey period by adding counts of sold and unsold chickens. Then, we divided this number by the number of days during which the same vendor bought any chickens, yielding an estimate of the number of chickens bought daily. We used these raw values to construct a discrete distribution from which to sample vendor batch sizes *S*_*V*_ in the ABM (Fig. A3A).

##### Surplus chickens

First, we estimated the probability *p*_*empty*_ to sell the entire batch of chickens before a vendor buys another batch. This was computed as the ratio between the number of days with any unsold chickens and the number of days with a purchase.

Second, we modelled the counts *u*_*batch*_ of unsold chickens per batch, conditional on batch size *S*_*V*_ and on the event that the batch is not fully emptied (which happens with probability 1 − *p*_*empty*_). We consider a negative binomial distribution with mean *μ* = *ρ*_*unsold*_*S*_*V*_ and variance 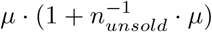, where *ρ*_*unsold*_ is the proportion of chickens that remain unsold before purchasing the next batch and *n*_*unsold*_ affects overdispersion compared to a Poisson distribution. Because our data are aggregated over a week period, we model the total count of unsold chickens *u*_*total*_ rather than *u*_*batch*_. Because the sum of negative binomial random variables still follows a negative binomial distribution, *u*_*total*_ has the same distribution as *u*_*batch*_, with *S*_*V*_ multiplied by the number of days with any unsold chickens (as we are conditioning on having a surplus). Aggregating data from wholesalers and retailers yields maximum likelihood estimates 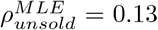 and 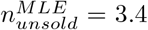 (Fig. A3B-C), which are used throughout the main manuscript. For completeness, we also estimate the same parameters for wholesalers and retailers separately, who amount to 55 and 376 data points, respectively. We find that wholesalers generate, on average and conditional on at least one unsold chicken, less surplus chickens than retailers 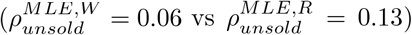, while the amount of overdispersion is roughly the same 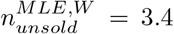 and 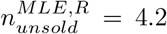). Note that we assume *p*_*empty*_ to differ between retailers and wholesalers. It should also be added that in the context of questionnaire data, a surplus of chickens is defined in relation to consecutive purchases, which could be multiple days apart, by the same vendor. In the ABM, however, we conflate these parameters with surplus between consecutive days, as vendors tend to buy new chickens every day. The resulting discrepancy should not be large since 76% of interviewed vendors purchased (broiler) chickens every day, and 93% during at least 6 days in a week. Finally, note that during simulations, daily surplus is also subject to the constrain that it can not be larger than batch size.

##### Prioritising unsold chickens

We estimate the probability of a vendor prioritising the sale of previously unsold chickens as the proportion of vendors declaring to do so in our data.

##### Transaction networks and inter-tier fluxes

Here we detail a procedure to estimate 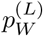 and 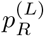 for each market tier *L* = 0, 1, …, *L*_*max*_. Parameters 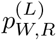 represent the proportions of chickens sold respectively to wholesalers and retailers in tier *L*, while 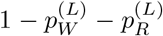 is the proportion of chickens that wholesalers in tier *L −*1 (*L >* 1) sell to end-point consumers.

The tier *L* = 0 corresponds to vendors that buy directly from middlemen. Note that 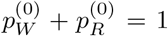 since middlemen are not allowed to sell chickens to end-point consumers. Wholesalers in tier *L* sell to wholesalers and retailers in tier *L* + 1 according to probabilities 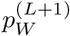 and 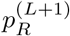, respectively. Note that the last tier *L*_*max*_ must consist of retailers only and hence 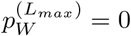.

Here we outline a simplified estimation procedure that yields parameters that are common to all markets. We start with reconstructed transaction networks. These networks are DAGs consisting of multiple transactions, weighted by the number *g* of chickens involved, between different node types. Nodes can be either middlemen (MM) or vendors, the latter being further classified into either wholesalers (W) or retailers (R). In addition, MM and W nodes are further classified according to their ‘depth’ within a transaction chain. This latter characterization is somewhat reminiscent of our tiered structure, but a direct mapping is not possible at this stage yet. In particular, in order to infer relevant parameters from reconstructed transaction chains, we must take care of transactions involving middlemen that purchase chickens directly from wholesalers and sell them to other vendors. As our ABM does not allow middlemen to buy from vendors, we replace these ‘forbidden’ transactions found in reconstructed networks, with sets of ‘allowed’ transactions. Let 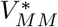 denote the set of such middlemen. As a first step, we remove all transactions involving poultry farms, except those involving any middlemen *k* ∈ 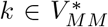. For each remaining farm *i*, we replace middlemen 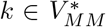 with a new label *k*^*∗*^ so that each transaction *i* → *k* 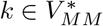 becomes *i* → *k*^*∗*^. The label *k*^*∗*^ is just a placeholder representing a fictitious middleman. Second, We then remove transactions of the type *i* → *k* and *k* → *j*, where *i, j* are not farms and 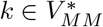, and replace them with viable transactions of the type *i →j*. Thus, if actor *i* sold chickens to 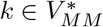, which in turn sold chickens to vendor *j*, we remove edges (*i, k*) and (*k, j*), and create a new edge (*i, j*) with a weight *g*_*i,j*_ = *g*_*i,k*_*g*_*k,j*_*/g*_*k*_, where *g*_*k*_ = Σ _*l*_ *g*_*k,l*_ is the total out-weight of *k*. Finally, we remove transactions involving only middlemen and merge any duplicated edges together.

We can now analyse the resulting graph *G*. Let *V*_*MM*_ represent the set of middlemen nodes, i.e. the roots of the DAG. Note that *V*_*MM*_ excludes nodes from 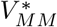. We associate middleman *i* with a weight 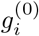 representing the number of chickens it ‘injects’ into markets. For each middleman node *i* ∈*V*_*MM*_, we then perform the following calculation:

- Enumerate all chains departing from node *i* and ending with a leaf node. The latter might be a W or a R node. We stress that these are observed transactions, not the output of our ABM.
- For each such chain, count the number of mark-ups, distinguishing between R and W nodes. Each chain may involve any number of W nodes and an optional terminal R node. For example, a chain of the type *W →W →R* consists of two wholesalers mark-ups before landing into a retailer. A chain of the type *W →W* consists of two wholesalers mark-ups, with the terminal wholesaler selling directly to end-consumers. Finally, assign a weight to the current chain by multiplying 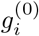 by the proportion of chickens that reach end-consumers through this chain. The latter is easily computed by multiplying together the proportions of chickens flowing through each edge in the chain. It is easy to see that the weights of chains emanating from node *i* must add up to 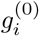.

By enumerating all chains constructed in this way, we can easily determine the maximum number of tiers as the length of the longest chain in *G*. For example, if the longest chain is of the type *W* → *W* → *R*, then we need 3 tiers and *L* = 0, 1, *L*_*max*_ = 2.

Once we have computed the weights for every chain in *G*, we can finally obtain 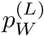 sand 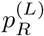 by collapsing all chains according to their length and the type of terminal node. The idea is to compute a running sum *g*_*G*_(*L, T*) that counts how many chickens end up in the hands of actors of type T, where T is either W, R or C (end-point consumer) in tier *L*. Please note that we are deliberately over-counting chickens according to how many mark-ups they make. Let us illustrate this procedure by considering a generic chain with weight *g*_*c*_, *n*_*W*_ *>*= 0 wholesaler and *n*_*R*_ = 0, 1 retailer mark-ups. The total length of the chain is *n*_*c*_ = *n*_*W*_ + *n*_*R*_. For each *L* = 0, …, *n*_*W*_ − 1 we increase the running sum *g*_*G*_(*L, W*) by *g*_*c*_. If the last node is of type W, we increase the running sum *g*_*G*_(*n*_*W*_, *C*) by *g*_*c*_; else, the last node is of type *R* and we increase the running sum *g*_*G*_(*n*_*W*_, *R*) by *g*_*c*_.

Finally, 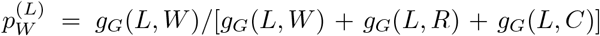 and 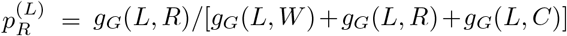. Note that *g*_*G*_(0, *C*) = 0 if we ignore transactions involving MM selling to end-point consumers.

Note that we can include more reconstructed transaction networks in the same analysis: we compute *g*_*G*_(*L, T*) ∀*G* and then create a consensus quantity *g*_*consensus*_(*L, T*) = Σ_*G*_ *g*_*G*_(*L, T*), which we can finally use to compute parameters of interest.

### 2 Actor dynamics

This section describes in detail different action and tasks for each actor. Further details on actor instantiation can be found in the section dedicated to PDN setup.

#### 2.1 Farms

At any time, a farm is either empty or raising chickens. We consider farms raising a single batch of chickens at a time, meaning that individual production cycles do not overlap. Chickens from the same batch are introduced at the same time as day-old chicks and are offered for sale as soon as they reach an appropriate age *τ*_*raise*_. A production cycle ends when all chickens in a farm are sold.

##### Farm replenishment

After completing a cycle, a farm remains empty for a random time *τ*_*replenish*_, sampled from a (shifted) negative binomial distribution. After *τ*_*replenish*_ time steps, a farm recruits a new batch of day-old chicks.

**Figure A 3:**
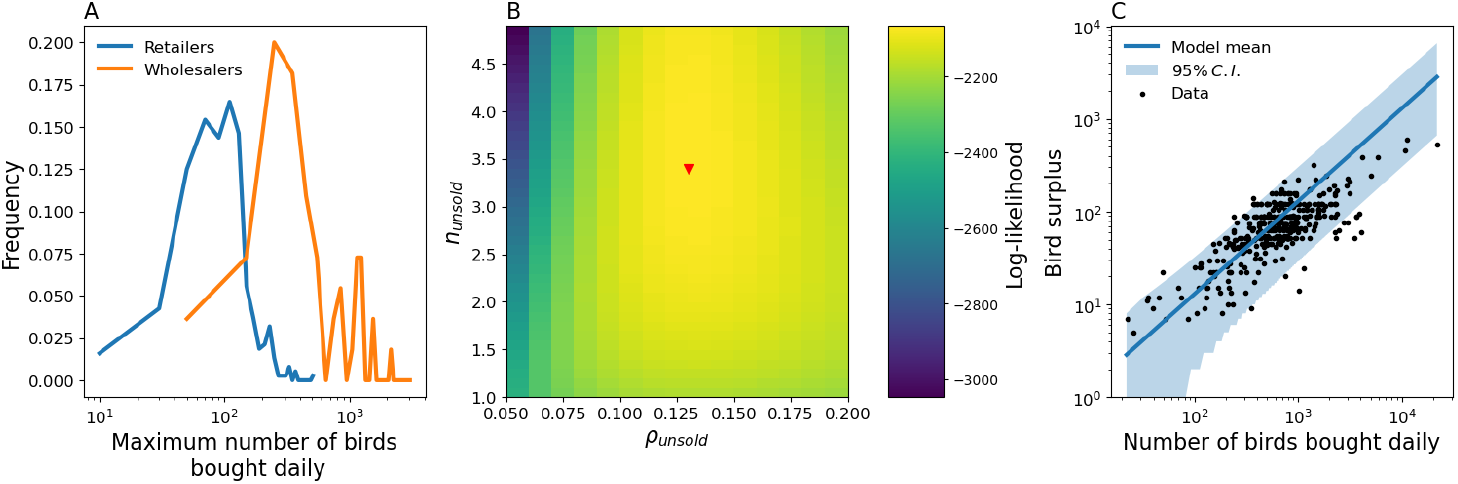
Vendors’ parameters analysis. (A) Raw distributions of number of birds bought daily for R (blue) and W (orange). (B) Log-likelihood function for negative binomial fit to surplus counts. Fitted parameters include the average proportion on unsold chickens *ρ*_*unsold*_ and the shape parameter *n*_*unsold*_. The maximum likelihood solution 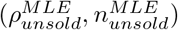 is indicated with a red marker. (C) Expected surplus 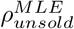 · *S* (line) as a function of batch size *S*, together with 95% C.I. (shaded area) calculated assuming a negative binomial distribution 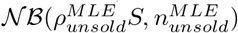 for surplus. Broiler data are shown as well (black dots).

##### Offering birds for sale

After raising a batch for *τ*_*raise*_ days, a farm is ready to sell chickens. These are offered for sale progressively over a minimum of *τ*_*rollout*_ days; more precisely, a farm stages new chickens corresponding to a fraction 1*/τ*_*rollout*_ of its batch over the first *τ*_*rollout*_ days. By day *τ*_*rollout*_, all chickens at the farm can be sold to middlemen. Note that a farm could sell chickens for a longer period of time if not enough middlemen are available to trade with. In the case where any chickens remain unsold after 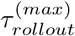 days, the corresponding farm is emptied automatically.

#### 2.2 Middlemen

Middlemen daily routine consists of buying and collecting chickens from farms in order to sell them to vendors operating in markets. A single middleman is able to source chickens from multiple farms, potentially from distinct regions, in order to fill up his/her cargo.

##### Updating scouted regions

Each middleman purchases chickens from farms located in a subset 𝒜 of neighboring areas, whose number we denote with *n*_*scout*_ = | 𝒜|. With daily frequency, each middleman may move and change his/her catchment area with probability *P*_*move*_. In that case, a middleman updates 𝒜by choosing *n*_*scout*_ new regions as follows: first, a focal region *a* is sampled at random with propensity proportional to the number of chickens being currently offered for sale in that region. In the case no farm is currently trading, *a* is chosen fully at random. Second, *n*_*scout*_ − 1 new regions are chosen at random among those neighboring *a*. More precisely, we say that a region *a*^*′*^ neighbors *a* if their centroids are less than 80 km far apart. If *a* has less than *n*_*scout*_ *−*1 neighbors, new regions are chosen among the areas neighboring with *a*’s direct neighbors. The choice of a 80 km radius is arbitrary, but it guarantees that middlemen do not cover too long distances in a single day. Note also that according to our algorithm, middlemen are drawn preferentially to areas with the largest offer of chickens, which reduces the odds of farms not being able to sell their chickens.

##### Buying birds from farms

The main purpose of mobile traders is to collect chickens from farms and deliver them to LBMs. With daily frequency, our algorithm allocates amounts of chickens to be moved from any trading farm to any market via middlemen. No chicken is collected at this stage yet. Our simulator resolves allocations one middleman at a time in random order and under the following constraints:

1. Middleman *j* can hold at most *S*_*MM*_ chickens.
2. Middleman *j* can source chickens from farms located in any area *a* ∈ A.
3. Middleman *j* sells chickens to a number of vendors from exactly *k*_*m*_ markets.
4. Destination markets must be chosen in a way the preserves the overall proportion of chickens leaving region *a* and entering market *l, f*_*a,l*_.

As a consequence of the last two requirements, each middleman keeps track of which market each chicken is scheduled to be moved to.

The first step consists in sampling markets with probabilities {*q*_*l*_}*l*=1,…, *N* _*M*_ :

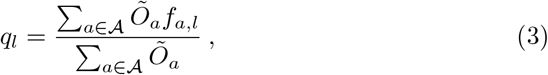

where *Õ*_*a*_ is the number of birds currently offered for sale by farms in region *a*. We sample markets with replacement until we get a set ℳ comprising *k*_*m*_ distinct markets, or after 1000 draws, in which case |ℳ| *< k*_*m*_. Let *t*_*l*_ be the number of times market *l* ∈ ℳ was selected during sampling, and let *t* = Σ _*l*_ *t*_*l*_ be the total number of draws. Focusing on market *l* with *t*_*l*_ *>* 0, we construct the distribution *f*_*a*|*l*_:

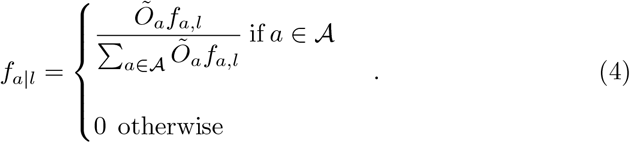

We then sample tokens *t*_*l,a*_ ∼ *Multinomial*(*t*_*l*_, {*f*_*a*|*l*_}) such that Σ _*a∈A*_ *t*_*l,a*_ = *t*_*l*_. Given the tokens *t*_*l,a*_, middleman *j* contacts farms in each region *a* ∈ A, securing up to 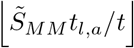 birds destined to market *l*. Here, 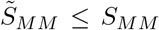 denotes the maximum number of birds that can be purchased during the current day. 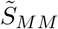 is smaller than *S*_*MM*_ if middleman *j* is already carrying birds for any reason, e.g. if unsold from previous days (this is usually rare).

Middlemen contact farms in an area *a* sequentially, starting with those trading for the longest amount of time. Transactions follow a greedy heuristics: if middleman *j* has to collect *s*_*a*_ birds from region *a*, he/she will attempt to buy as many birds as possible from individual farms to meet such quota. In case middleman *j* fails to secure the required number of birds from region *a*, we allow *j* to contact any previously visited farms to meet the original quota. Additional chickens obtained in this way are directed to market *l* with probability:

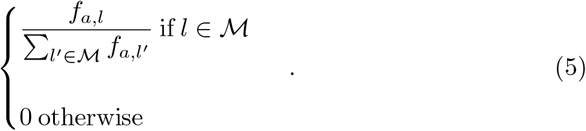

Finally, it should be noted that *Õ* _*a*_ decreases as chickens are allocated progressively to middlemen; in other words, availability of chickens decreases as we iterate over middlemen.

##### Bird collection

Middlemen start collecting birds as soon as the allocation step described in the previous section is terminated. At this point, middleman *j* is already aware of the farms to be visited and the amounts of birds to pick up from each of them. In simulations, middleman *j* can visit any of these farms in random order at any point in time before the next market opening. This is to ensure that *j* collects all of their chickens before bringing them to the market.

##### Selling birds to vendors

Each day, middleman *j* visits markets *l* ∈ M sequentially. There, *j* sells chickens to available vendors until all *s*_*l*_ carried chickens scheduled for delivery to market *l* have been sold. These chickens can be sold only to vendors in the first market tier (*L* = 0). A number 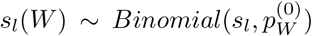 are allocated for sale to wholesalers in this tier. The remaining *s*_*l*_ −*s*_*l*_(*W*) chickens, plus any of the *s*_*l*_(*W*) chickens that could not be sold to wholesalers (e.g. because of limited buying capacity) are directed to retailers. Any unsold chickens remain in middleman’s stock and are offered for sale in the same market on the following day. Please note that this event is expected to occur rarely in simulations.

#### 2.3 Markets

Markets can be either open or closed: all markets open at *T*_*open*_ and close at *T*_*close*_ every day. Markets contain chickens only during opening hours.

#### 2.4 Vendors

Vendors trade chickens within and between markets. There are two types of vendors: retailers, who sell chickens to end-point consumers only, and wholesalers, who can also sell chickens to other vendors. Vendors are organized in tiers: vendors in tier *L >* 0 buy chickens from wholesalers in tier *L* −1, while vendors in tier *L* = 0 buy chickens directly from middlemen. Vendors first purchase chickens in market *l* and then move to another market *l*^*′*^, or stay in *l*, where they can sell chickens to other vendors and/or end-point consumers.

##### Buying birds

All vendors seek to buy as many chickens as possible during any transaction with middlemen and/or other vendors, compatibly with their own capacity and daily quota *S*_*V*_. Note that a vendor may hold more than *S*_*V*_ chickens at a time due to the presence of surplus chickens. Nonetheless, the same vendor can not hold more chickens than the maximum carrying capacity *C*_*V*_ *> S*_*V*_.

##### Inter-market movements

At *T*_*open*_, vendor *k* moves to market *l* to purchase chickens. Eventually, *k* stays in the same market or moves to a second location *l*^*′*^ to trade. Chickens move alongside their owner. Note that *k*’s purchase and trade markets *l,l*^*′*^ are invariant. In other words, *k* will always purchase chickens in *l* and commute to *l*^*′*^ (if *l*≠ *l*^*′*^) or stay in *l* (if *l* = *l*^*′*^) in a given PDN realisation.

All vendor movements are resolved at *T*_*open*_ and are therefore instantaneous: vendors move to market *l*, purchase chickens and eventually change market during the same time step. Note that vendors in tier *L* must move after those in tier *L* − 1.

##### Wholesaling

Let us consider a wholesaler *k* operating in tier *L*. After purchasing chickens, *k* carries *n*_*tot*_ chickens, including also older chickens that remained unsold from previous days. In the current day, *k* sells *n*_*sale*_ = *n*_*tot*_ − *n*_*unsold*_ chickens, where *n*_*unsold*_ denotes the number of unsold chickens. *n*_*unsold*_ is a random variable whose sampling procedure is described in sections below. Proportions 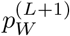 and 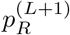 of these *n*_*sale*_ are then directed to wholesalers and retailers in tier *L* + 1, respectively. The remaining chickens are sold to end-point consumers (retailing).

It is possible that *k* sells less chickens to *L* + 1 tier wholesalers than planned. In this case, any surplus chickens are redirected to retailers. Similarly, any chickens that would remain unsold after the wholesaling phase are redirected to end-point consumers.

Please note that wholesaling is instantaneous, occurring toe-to-toe with inter-market movements as vendor switch from purchasing to trading chickens.

##### Retailing

Both wholesalers and retailers can sell chickens to end-point customers. While wholesaling is instantaneous, retailing rolls out over market opening hours, i.e. between *T*_*open*_ and *T*_*close*_. More in detail, vendors can sell chickens to end-point consumers between *T*_*open*_ + 1 and *T*_*close*_ −1; this means that chickens spend at least one time step at the market.

The number of chickens sold in any time step over this period is stochastic, but uniformly distributed. More in detail, we assume that the number of chickens sold by vendor *k* during time step *t* is given by:

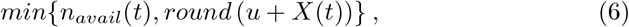

where *X* is a Poisson random variate with mean *n*_*avail*_(*t*)*/δt*, with *n*_*avail*_(*t*) and *δt* denoting the number of chickens destined to retail still in stock and the time left until market closes, respectively. *u* is a random draw from an uniform distribution on the unit interval whose role is to fix rounding issues, while *round*(.) guarantees a meaningful result by rounding *u* + *X* to the nearest integer. Finally, we require that *k* sells all of the remaining chickens that had been scheduled for retail, right before market closure, i.e. at *t* = *T*_*close*_ − 1.

##### Surplus chickens

As explained above, vendor *k* may retain a random number *n*_*unsold*_ of chickens each day, which are then offered for sale the day after. The surplus *n*_*unsold*_ is computed as follows: first, vendor *k* sells the entire stock with probability *p*_*empty*_, in which case *n*_*unsold*_ = 0. Alternatively, *n*_*unsold*_ is sampled from a negative binomial distribution with mean *ρ*_*unsold*_ · *n*_*tot*_, with the constraint *n*_*unsold*_ *≤n*_*tot*_. More details can be found in the data analysis section.

##### Prioritising unsold birds

Vendors keep track of unsold birds. We assume that a proportion *P*_*priority*_ of vendors in our simulations prioritise selling these birds before recently purchased birds. The remaining vendors, instead, do not prioritise repurposed chickens over those bought in the current day.

### 3 PDN setup

This section illustrates the generative algorithm responsible for instantiating the PDN. The following subsections reflect the sequential steps of the algorithm. The final goal is to generate a kind of supply chain with multiple intermediate nodes under certain structural constraints.

The algorithm first instantiates ‘source’ and ‘sink’ nodes, i.e. farms and markets, where chickens are first introduced and sold last, respectively. Then, it populates markets with vendors, compatibly with incoming bird flux as determined by farm production and region-to-market fluxes. Finally, our algorithm instantiates middlemen, i.e. the intermediate actors.

We assume here for simplicity that all actors handle and trade a single chicken breed.

#### 3.1 Geography setup

Using external specifications about the study area, the algorithm creates a list of *N*_*A*_ regions, each being characterized by a given location and a weight proportional to the share 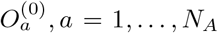 of chickens produced there. In the case of Bangladesh we identify individual regions with upazilas. In addition, we also specify a matrix *f*_*a,l*_ representing the proportion of chickens that end up in market *l* = 1, …, *N*_*M*_ from area *a*. Note that Σ_*l*_ *f*_*a,l*_ = 1.

#### 3.2 Farm generation

##### Farm properties

In this work we consider only farms raising a single batch of chickens at a time. For this type of farm then, farm size *S*_*F*_ is equivalent to batch size; the latter is assumed to vary across farms and is sampled from a truncated negative binomial distribution. In addition, batch size is invariant, in the sense that a particular farm will always raise batches with the same size. The raising time *τ*_*raise*_ is assumed to be a constant and shared by all farms. Analogously to batch size, a random value of minimum rollout duration *τ*_*rollout*_ is assigned to each farm according to a probabilistic distributions but does not change over the course of a simulation. Finally, replenishment time *τ*_*replenish*_ is assumed to be sampled from a shifted negative binomial distribution, namely 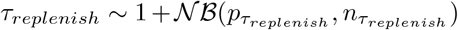, whenever a farm completes a rollout. Farm properties and model parameters describing farm generation are listed in Table A1.

##### Farm locations

For each farm we draw a random location according to the following algorithm: we first draw a region *a*, either uniformly at random with probability *P*_*random*_, or proportionally to the outgoing flux 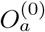 with complementary probability. Then, we sample a random point within the selected region.

Alternatively, empirical or simulated data on farm spatial distributions could be used to fix their locations.

##### Farm output

Once all farms have been generated, we compute a range of quantities that are fundamental to PDN generation and/or dynamics.

First, we compute the expected daily bird output *O*_*i*_ from farm *i* = 1, …, *N*_*F*_. We compute *O*_*i*_ by assuming that farm *i* completes every rollout in exactly *τ*_*rollout,i*_ days yielding:

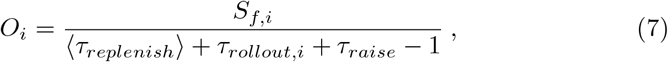

it should be noted, however, that this overestimates the true expected daily output *O*_*i*_ since *τ*_*rollout,i*_ represents only the minimum rollout duration, which might take longer in absence of middlemen to collect birds. Nonetheless, we will stick to the heuristic calculation in eq. S(7) to evaluate other quantities. These include expected daily bird output from region *a*:

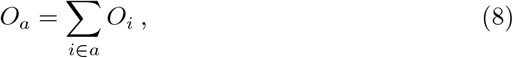

where the sum runs over all farms located in region *a*. Given the regional outputs, it is possible to compute the following quantities:

- *Õ* _*a*_ = *O*_*a*_*/* Σ _*a*_^*′*^ *O*_*a*_^*′*^, a normalized version of *O*_*a*_.
- *q*_*a,l*_ = *O*_*a*_ · *f*_*a,l*_, the expected daily bird flux from region *a* to market *l*.
- *M*_*l*_ = Σ _*a*_ *q*_*a,l*_, the expected daily bird flux impinging on market *l*.

**Table A 1:** Farm-specific parameters. A fraction *f*_*random*_ of farms are allocated in random upazilas, while remaining farms are assigned to upazilas proportionally to their volume of traded chickens. Farm locations are completely random within upazilas. Farm size and replenishment time distributions are 𝒩 ℬ (*p, n*). Farm size *S*_*F*_ is further constrained in the range 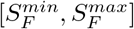. Farm size is sampled once per farm per realisation: a farm always recruits the same amount of birds. Refill and rollout times are sampled during each production cycle.

| Parameter | Meaning | Distribution/Value | Source |
| --- | --- | --- | --- |
| $N_F$ | Number of farms | 1200 | - |
| $f_{random}$ | Prob. random allocation | 0.3 | Assumed |
| $T'$ | Farm update time | 11 a.m. | - |
| $S_F$ | Farm size | $\mathcal{NB}(p_{S_F}, n_{S_F})$ | Estimated |
| $p_{S_F}$ | Farm size $\mathcal{NB}$ parameter | 0.003544 | Estimated |
| $n_{S_F}$ | Farm size $\mathcal{NB}$ parameter | 5.245 | Estimated |
| $S_F^{min/max}$ | Min/max farm size | 600, 4180 | Estimated |
| $\tau_{raise}$ | Raising time | 32 days | [1] |
| $p_{\tau_{replenish}}$ | $\tau_{replenish}$ $\mathcal{NB}$ parameter | 0.2013 | Estimated |
| $n_{\tau_{replenish}}$ | $\tau_{replenish}$ $\mathcal{NB}$ parameter | 4.488 | Estimated |
| $P(\tau_{rollout})$ | Min. rollout time distribution | (0.2, 0.3, 0.3, 0.2)<br>for $\tau_{rollout} = 1 - 4$ days | Assumed |

#### 3.3 Market setup

Details about markets are provided externally. At this stage, the algorithm instantiates *N*_*M*_ markets, each structured in *L*_*max*_ + 1 empty tiers. Market properties and model parameters describing market generation are listed in Table A2.

**Table A 2:** Market parameters.

| Parameter | Meaning | Distribution/Value | Source |
| --- | --- | --- | --- |
| $N_M$ | Number of LBMs | 20 | - |
| $T_{open}$ | LBM opening time | 6 a.m. | - |
| $T_{close}$ | LBM closing time | 11 p.m. | - |
| $w_{l,l'}$ | Prob. vendor purchasing in LBMs $l$ sells in $l'$ | Explored | Modelled |
| $f_{a,l}$ | Prop. of chickens leaving upazila $a$ for LBM $l$ | Empirical | [2] |

#### 3.2 Vendor setup

##### Vendor properties

At setup, each vendor is assigned a batch size *S*_*V*_, drawn from a probabilistic distribution, the latter being different for W and R. Here, *S*_*V*_ denotes the maximum number of chickens he/she can purchase in a single day. Parameters describing properties of vendors and their generation are listed in Table A3, while parameters relating to market tiers are listed in Table A4.

##### Tier-by-tier vendor allocation

As explained in the main text, each vendor operates in up to two markets, and always in the same tier. Our algorithm assigns vendors to markets and tiers within them in a way that is compatible with the expected flux of chickens entering the market, as well as with between-tier fluxes. Importantly, neither the number of W and R can be specified a priori as they are determined by our algorithm, which we define as follows.

Starting from tier *L* = 0, we assign as many R and W to that tier so that their combined capacities *S*_*V*_ match the expected R and W incoming fluxes 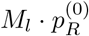 and 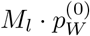 from middlemen. Then, we assign a random destination market *l*^*′*^ with probability *w*_*l,l*_^*′*^ to each vendor generated in the current tier; If *l*^*′*^ = *l*, a vendor will operate in a single market.

We then proceed by evaluating the expected flux of chickens 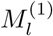 impinging on tier *L* = 1 due to vendors in tier *L* = 0; this amounts to compute the combined expected output from all *L* = 0 wholesalers buying chickens in any market and selling in *l*. The expected wholesaler output is taken to be equal to *S*_*V*_. Given 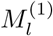, we allocate as many R and W to match fluxes 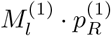 and 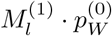. Finally, we repeat the same scheme for the deeper tiers as well.

As a final note, we allow for a multiplicative scaling factor, to be applied to 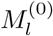, whose effect is to allocate more vendors than what would be implied by our heuristic calculation of farm outputs.

Note that a vendor buying *S*_*V*_ chickens daily also sells the same amount of chickens on average. To see that, let us consider the following discrete-time process: during time step *t*, a vendor accumulates *S*_*V*_ chickens and sells all of his stock with probability *p*_*empty*_, or a proportion 1 − *ρ*_*unsold*_ with complementary probability. The average surplus at time step *t* + 1 is given by:

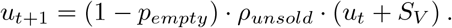

At stationariety we must have *u*_*t*+1_ = *u*_*t*_ *≡u*^*∗*^, which implies that the amounts of chickens sold and purchased must balance each other, hence our claim. Solving for *u*^*∗*^ yields:

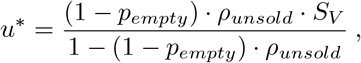

and an average daily stock, after acquiring *S*_*V*_ chickens:

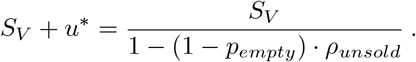

Finally, we take an individual’s vendor maximum carrying capacity *C*_*V*_ as being 1.5 times the average daily stock:

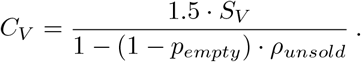

**Table A 3:** Vendor-specific parameters. W,R denote wholesalers and retailers, respectively. When some surplus is generated (with probability *p*_*empty*_), it is sampled from a negative binomial distribution with mean *m* = *ρ*_*unsold*_ *·n*, where *n* denotes total chickens offered for sale, and overdispersion parameter *α*_*unsold*_ (the variance is *m* + *α*_*unsold*_ ∗ *m*^2^).

| Parameter | Meaning | Distribution/Value | Source |
| --- | --- | --- | --- |
| $S_V$ | Max chickens purchased | Empirical | [2] |
| $p_{empty}$ | Prob. no surplus | 0.32 (W), 0.15 (R) | Estimated |
| $\rho_{unsold}$ | Prop. surplus | 0.13 (W & R) | Estimated |
| $\alpha_{unsold}$ | Overdispersion surplus | 0.29 (W & R) | Estimated |
| $P_{priority}$ | Prop. vendors selling older chickens first | 0.84 (W), 0.49 (R) | Estimated |

#### 3.5 Middlemen setup

##### Middlemen properties

Middlemen are assumed to trade a single chicken breed. Each middleman is assigned a capacity *S*_*MM*_ from a discrete distribution, denoting the maximum number of chickens it can buy in a single day. Analogously to vendors, the number of middlemen can not be specified a priori, as it is determined dynamically: we allocate as many vendors so that their combined capacity matches the expected daily output from farms, i.e. Σ_*i*_ *O*_*i*_.

**Table A 4:** Market tiers’ parameters. Parameters 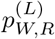 represent the proportions of chickens sold respectively to wholesalers and retailers in tier *L* from either middlemen (if *L* = 0) or wholesalers from the previous tier (if *L >* 0). 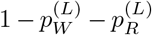 represents instead the proportion of chickens that wholesalers in tier *L* − 1 (*L >* 1) sell to end-point consumers (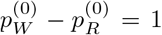 since middlemen do not sell to end-point consumers). Values are the same for all markets. Parameters are estimated from reconstructed transaction networks. The last tier contains only zeros as vendors in the previous tier sell only to end-point consumers.

| $L$ | $p_W^{(L)}$ | $p_R^{(L)}$ |
| --- | --- | --- |
| 0 | 0.606 | 0.394 |
| 1 | 0.407 | 0.315 |
| 2 | 0.169 | 0.318 |
| 3 | 0 | 0.058 |
| 4 | 0 | 0. |

In addition, each middleman is assigned an integer *k*_*m*_, sampled from a probability distribution, representing the number of markets visited daily. The middleman will then commit to sell chickens to *k*_*m*_ different markets during each day (*k*_*m*_ does not change in time). Properties of middlemen and model parameters describing middlemen generation are listed in Table A5.

##### Middlemen initial positions

As explained in the main text, at any point in time, each middleman tracks a set of *n*_*scout*_ distinct regions. During PDN generation, we assign middlemen to regions as follows: we first select an initial area *a* uniformly at random. Then, *n*_*scout*_ − 1 new areas are chosen among those neighboring *a*, i.e. within a distance *d < d*_*MM*_ (based on their centroids). In case not enough areas are selected, we choose among neighbors’ neighbors.

### 4 Simulating epidemic spread

Our simulator allows to simulate transmission of multiple pathogens/strains in the same poultry population. The current version of the simulator allows to simulate transmission across multiple scales, including at the level of the same flock and at the level of farms. All parameters describing pathogen transmission are listed in Table A6.

**Table A 5:** Middleman-specific parameters. Number of markets serviced daily is geometric with probability 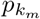 (truncated above by the number of markets). *k*_*m*_ is sampled only once per middleman per realisation.

| Parameter | Meaning | Distribution/Value | Source |
| --- | --- | --- | --- |
| $S_{MM}$ | Cargo size | Empirical | [2] |
| $P_{move}$ | Daily movement prob. | 0.1 | Assumed |
| $n_{scout}$ | Number of scouted areas | 4 | Assumed |
| $d_{MM}$ | distance between neighboring areas | 80 km | Assumed |
| $p_{k_m}$ | Markets serviced parameter | 0.66 | [2] |

#### 4.1 Within-flock transmission

Let us consider a population of chickens within a single setting, e.g. a farm, a middleman’s truck, a market, or a vendor’s shed. Note that at any time, any chicken belongs to one and only one setting.

We simulate transmission using Sellke’s construction [3]. Briefly, we assign a hazard value *h* to each chicken, sampled from an exponential distribution with unit rate; then, any contact with an infectious chicken reduces the target chicken’s hazard by an amount *δh*. Whenever *h* hits 0 due to an infectious contact, the target chicken is infected. Later, *h* is updated with another draw from an exponential distribution with unit rate.

We make the assumption that chickens within the same setting mix homogeneously at random. Therefore, infectious contacts are directed at random chickens. During simulations, each infectious chicken makes exactly one contact per time step with a randomly chosen chicken from the same setting.

Infection triggers a chain of events depending on the specified compartmental model. In SIR-like dynamics, an infected chicken becomes infectious immediately, but recovers after an infectious period 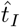 sampled from a geometric distribution with PMF:

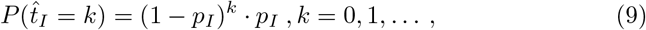

where *p*_*I*_ = 1 −*exp*(−(*T*_*I*_ + 1)^*−*1^) and *T*_*I*_ is the average infectious period. It is easy to check that the distribution in Eq. (9) has mean *T*_*I*_. In simulations, 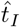 is sampled immediately after an infection happens, say time *t*, and recovery is deferred to time 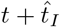. In models with latency, i.e. SEIR-like models, chickens become infectious only after an incubation time 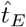 sampled from a geometric distribution with mean 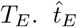 is sampled immediately after infection at time *t*, and the chicken becomes infectious only at time 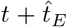, at which point the corresponding infectious period is also sampled.

Let us now consider an infectious chicken *i* trying to infect chicken *j* with pathogen *x*. The overall hazard reduction *δh* can then be written as:

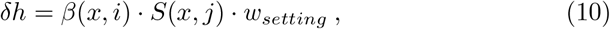

where *β*(*x, i*) is the transmissibility of *x* and *S*(*x, j*) is susceptibility of chicken *j* to infection with *x*. The factor *w*_*setting*_ is a multiplier that depends only on the current setting, and accounts for differences in transmission across settings. The factors *β*(*x, i*) and *S*(*x, j*) may depend on the state of the infector and infectee, respectively. Susceptibility may account for example for previous exposure to the same pathogen, or cross-reactions induced by exposure to other pathogens/strains. In the single-strain SIR model, for example, *S*(*x, j*) = 0 if *j* is infectious or recovered.

#### 4.2 Between-farm transmission

We allow pathogens to spread between distinct farms. If a farm *f* contains infectious chickens, the probability of infecting another farm *f*^*′*^ (provided *f*^*′*^ is not empty), irrespective of whether *f*^*′*^ is already infected or not, is computed as:

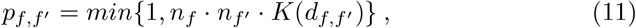

where *n*_*f*_, *n*_*f*_^*′*^ are the average numbers of chickens in farms *f* and *f*^*′*^, respectively, *d*_*f,f*_^*′*^ is the distance between *f* and *f*^*′*^ and *K*(*x*) is a spatial kernel. If transmission occurs, a random chicken in *f*^*′*^ is set as infected, conditional on not being already immune, and a random infector chicken is selected from farm

*f*. If the infector is co-infected with multiple strains, a single carried strain *s* is chosen at random and transmitted to the infectee. Note that this process bypasses the hazard rate calculation and that all strains are equivalent in the context of between-farm transmission.

We simulate transmission between farms using the Conditional Entry algorithm [4]. The algorithm requires farms to be assigned to cells in a grid in order to exploit the fact that transmission is more likely to occur within cells than between them. We use an adaptive algorithm described in the same paper to construct a grid over the farm population. The algorithm relies on a hyperparameter *λ*, here set to 15, that only affects the sizes and number of individual cells. Finally, because performing between-farm transmission is computationally expensive, we run the conditional entry algorithm only once per day rather than every time step.

In this work we consider a power-law transmission kernel:

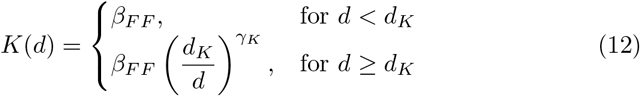

where *β*_*F F*_ denotes the overall strength of spatial transmission.

#### 4.3 External introductions

External transmission events are responsible for seeding and re-seeding pathogens in farms. Once a day we iterate over all farms and reduce the hazard of a randomly selected chicken *i* due to pathogen *s* by an amount:

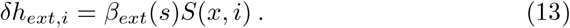

In the main manuscript, we also consider an alternative seeding protocol that introduces different strains in distinct upazilas.

**Table A 6:** Epidemic parameters. Distributions of latent and infectious period are geometric with expected values typical of AIV infections. The transmission kernel’s shape and parameters *γ*_*K*_ and *d*_*K*_ are instead inspired to a study of H5N1 epidemics in Dhaka region.

| Parameter | Meaning | Distribution/Value | Source |
| --- | --- | --- | --- |
| $T_E$ | Mean latent period | 6 hours | [5] |
| $T_I$ | Mean infectious period | 48 hours | [5] |
| $\beta$ | Base within-flock transmissibility | 0.2 hours<sup>-1</sup> | Assumed |
| $\beta_{ext}$ | Intro. rate | 0.0005 days<sup>-1</sup> | Assumed |
| $\beta_{FF}$ | Inter-flock transmissibility | $5 \cdot 10^{-11}$ days<sup>-1</sup> | Assumed |
| $\gamma_K$ | Exponent transmission kernel | 0.94 | [6] |
| $d_K$ | Scale transmission kernel | 0.1 km | [6] |
| $w_X$ | Setting-specific transmission multiplier | 0.2 (F), 1 (MM)<br>1 (M), 1 (V) | Assumed |

## Supplementary figures

**Supplementary Figure 1:**
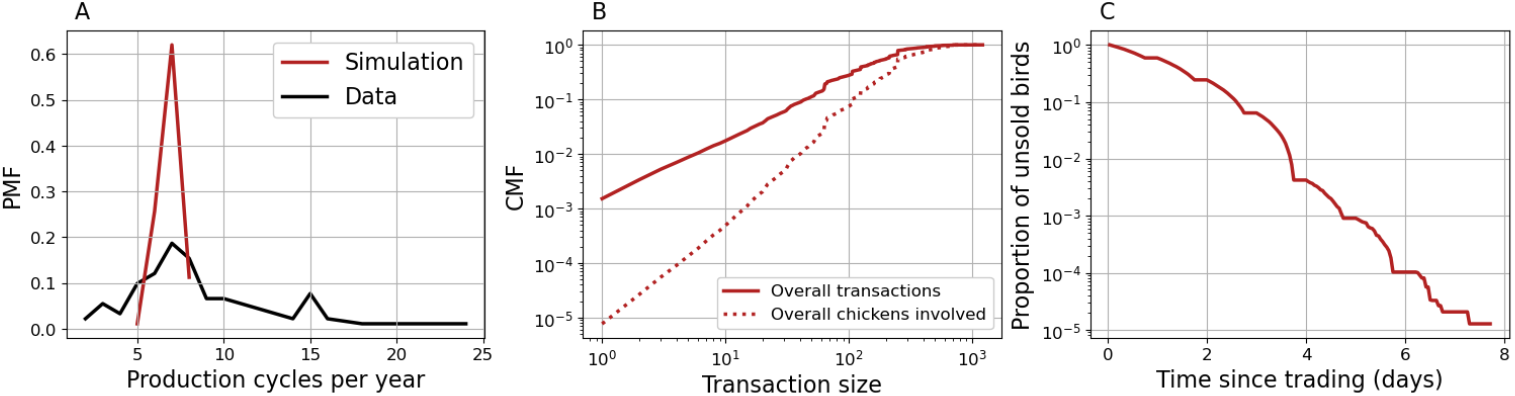
Additional farm statistics from simulations. (A) Distribution of numbers of production cycles completed per year. The simulated distribution (red) appears narrower compared to empirical data (black) [1]. However, it should be added that several interviewed farmers raised multiple batches simultaneously, and those that declared raising a single batch during the interview may well have being managing 2 or more simultaneously during the previous year. (B) Cumulative distribution of sizes of transactions involving farms and middlemen (solid line). The dotted line represents the cumulative proportion of chickens sold in transactions up to a given size. given size. The corresponding PMFs, denoted with *p*_*s*_ and *p*^*′*^_*s*_ respectively, are related since *p*^*′*^_*s*_ = *s· p*_*s*_ *Σ*_*s*_ *s· p*_*s*_. In other words, *p*^*′*^*s* is the size-biased version of *p*_*s*_. (C) Proportion of chickens remaining unsold after a given time since being offered for sale for the first time by a farmer. Note that it is highly unlikely for a chicken to remain unsold for more than 5 days. Results are obtained from a single simulation with default settings as in Fig. 2 in the main manuscript.

**Supplementary Figure 2:**
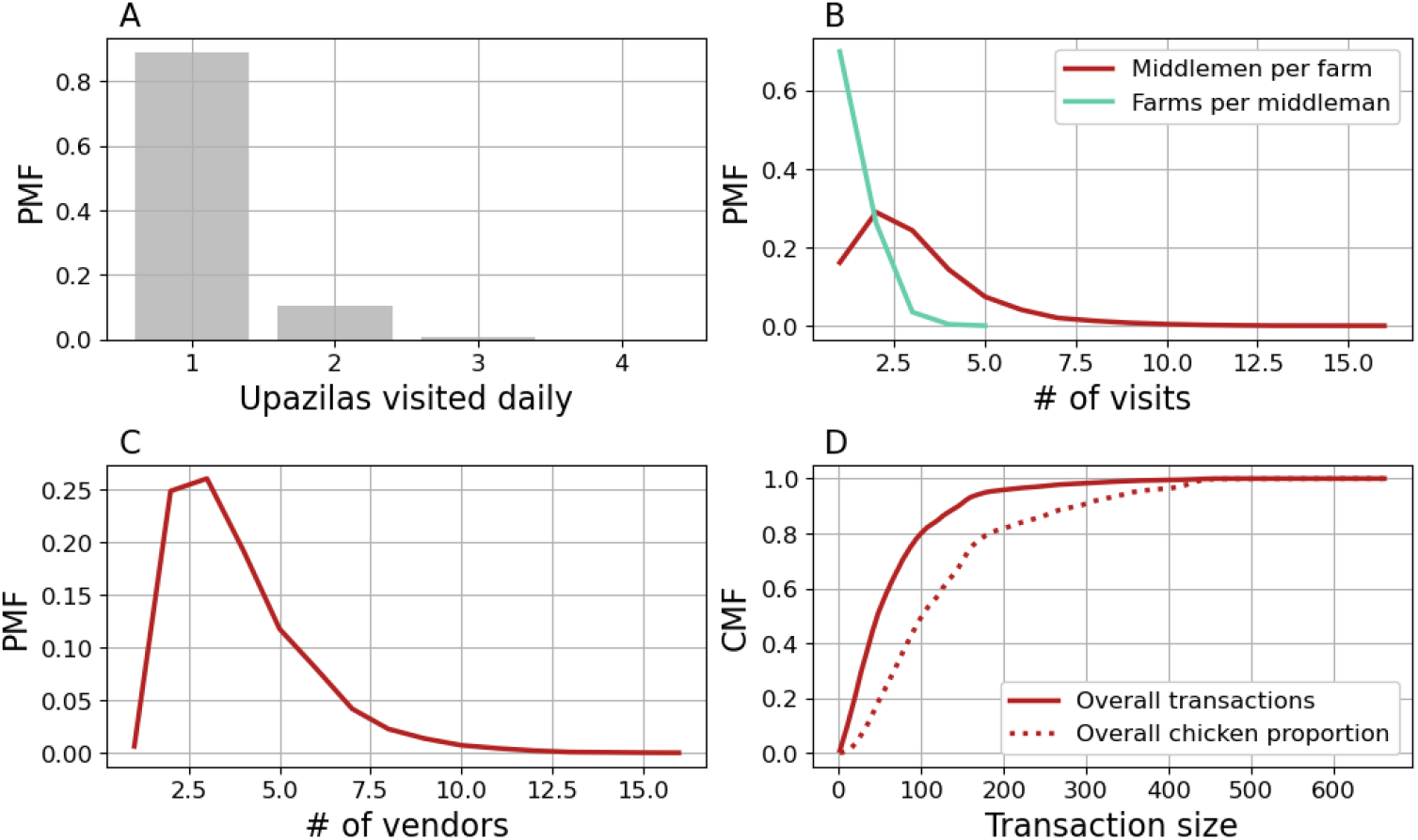
Additional middlemen statistics from simulations. (A) Proportion of upazilas visited daily by one middleman during a single simulation. Note that a middleman may visit up to 4 upazilas per day, but visiting one or two is usually sufficient to complete a cargo. (B) Distributions of daily numbers of farms visited by one middlemen (green) and middlemen visiting one farm (red). (C) Distribution of numbers of vendors trading daily with a middleman. (D) Cumulative distribution of sizes of transactions involving middlemen and vendors (solid line). The dotted line represents the cumulative proportion of chickens sold in transactions up to a given size. Note that these transactions are typically smaller than those between farms and middlemen since vendors deal with smaller amounts of chickens than other PDN actors. Results are obtained from a single simulation with default settings as in Fig. 2 in the main manuscript.

**Supplementary Figure 3:**
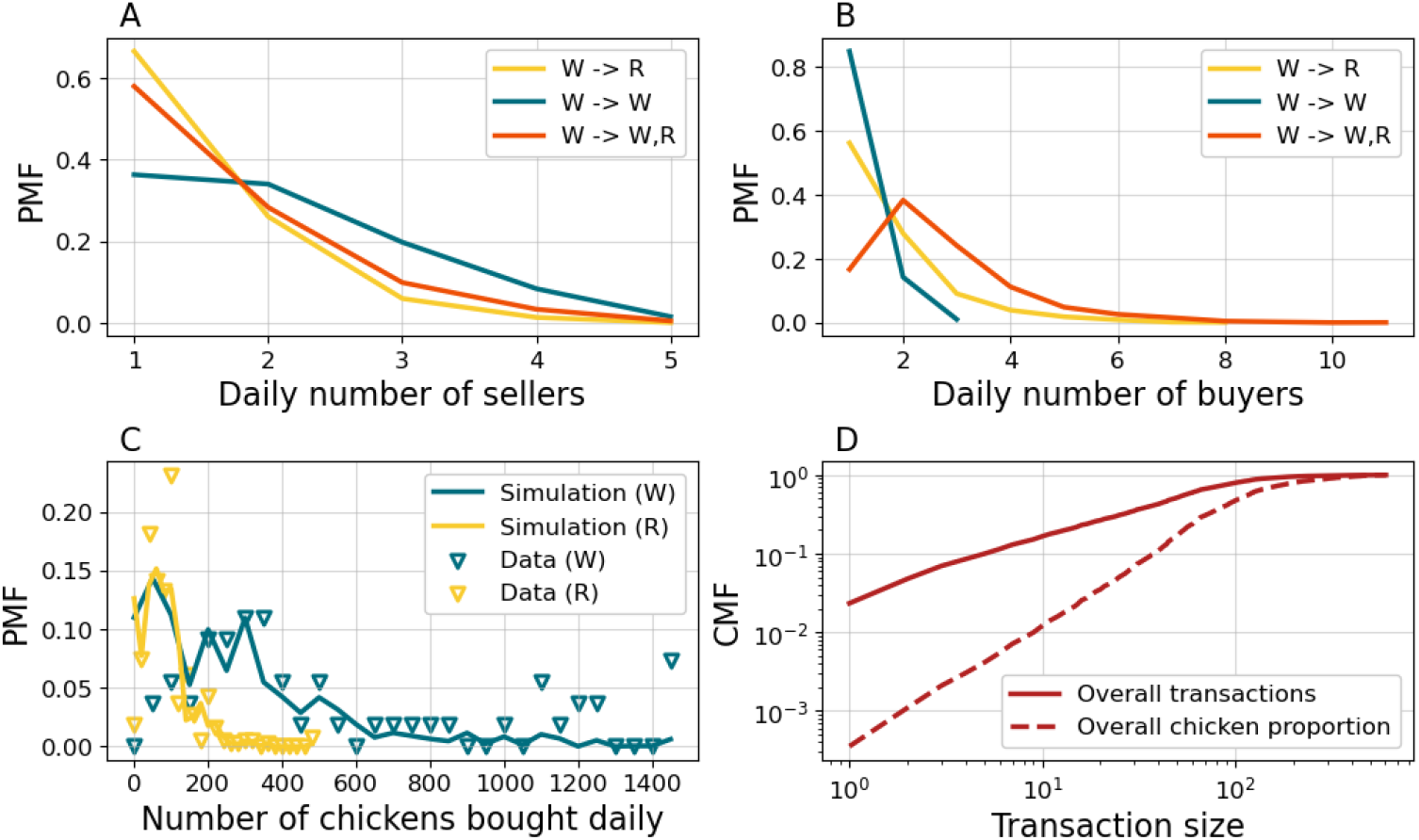
Additional vendor statistics from simulations. (A) Distribution of numbers of wholesalers supplying a retailer (yellow), another wholesaler (blue) or any vendor (red) on a daily basis. (B) Distribution of numbers of retailers (yellow), wholesalers (blue) or vendors (red), regardless of type, purchasing from a single wholesaler on a daily basis. Note that (A) excludes vendors buying chickens from middlemen, i.e. vendors operating in the first LBM tier. (C) Distributions of daily amounts of chickens bought from retailers (yellow) and wholesalers (blue) in simulations (lines) and data (markers) [2]. (D) Cumulative distribution of sizes of transactions involving middlemen and vendors (solid line). The dotted line represents the cumulative proportion of chickens sold in transactions up to a given size. Results are obtained from a single simulation with default settings as in Fig. 2 in the main manuscript.

**Supplementary Figure 4:**
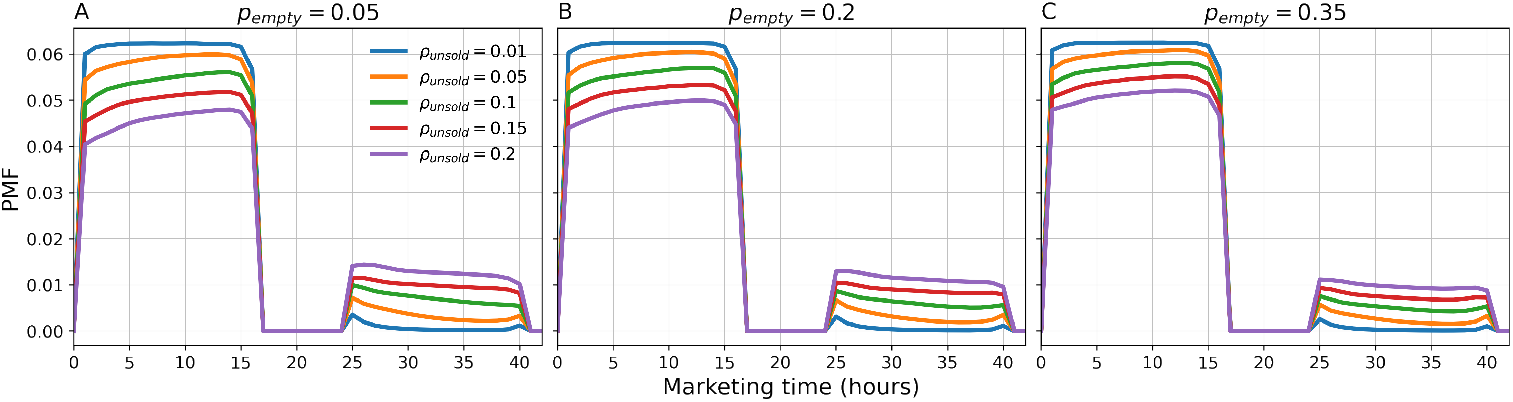
Distribution of marketing times. Each panel shows distributions of marketing times for different average proportions of unsold chickens *ρ*_*unsold*_ and for increasing probability *p*_*empty*_ of a vendor selling all chickens in a single day (from left to right). The marketing time is defined as the time interval elapsed since a chicken enters any LBM for the first time and is sold to an end-point customer. Simulation settings are the same as in Fig. 4 with only 10% of vendors prioritizing the sale of unsold chickens.

**Supplementary Figure 5:**
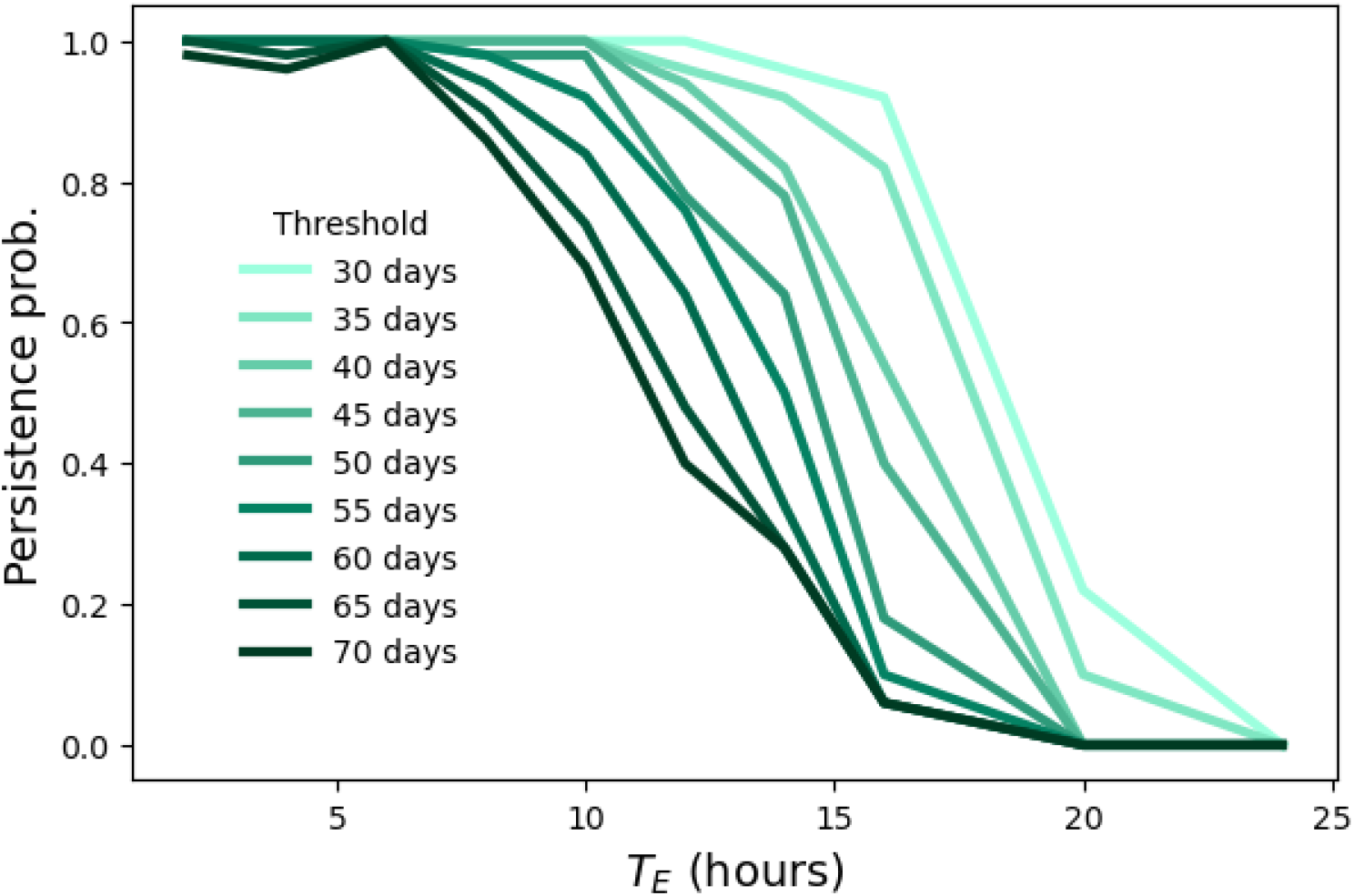
Sensitivity of persistence probability to duration of transmission chains. Lines show how the probability of pathogen persistence varies with both *T*_*E*_ and the minimum duration to determine whether a transmission chain is persistent or not. The estimation of the probability of persistence as well as simulation settings are the same as in Fig. 6I.

**Supplementary Figure 6:**
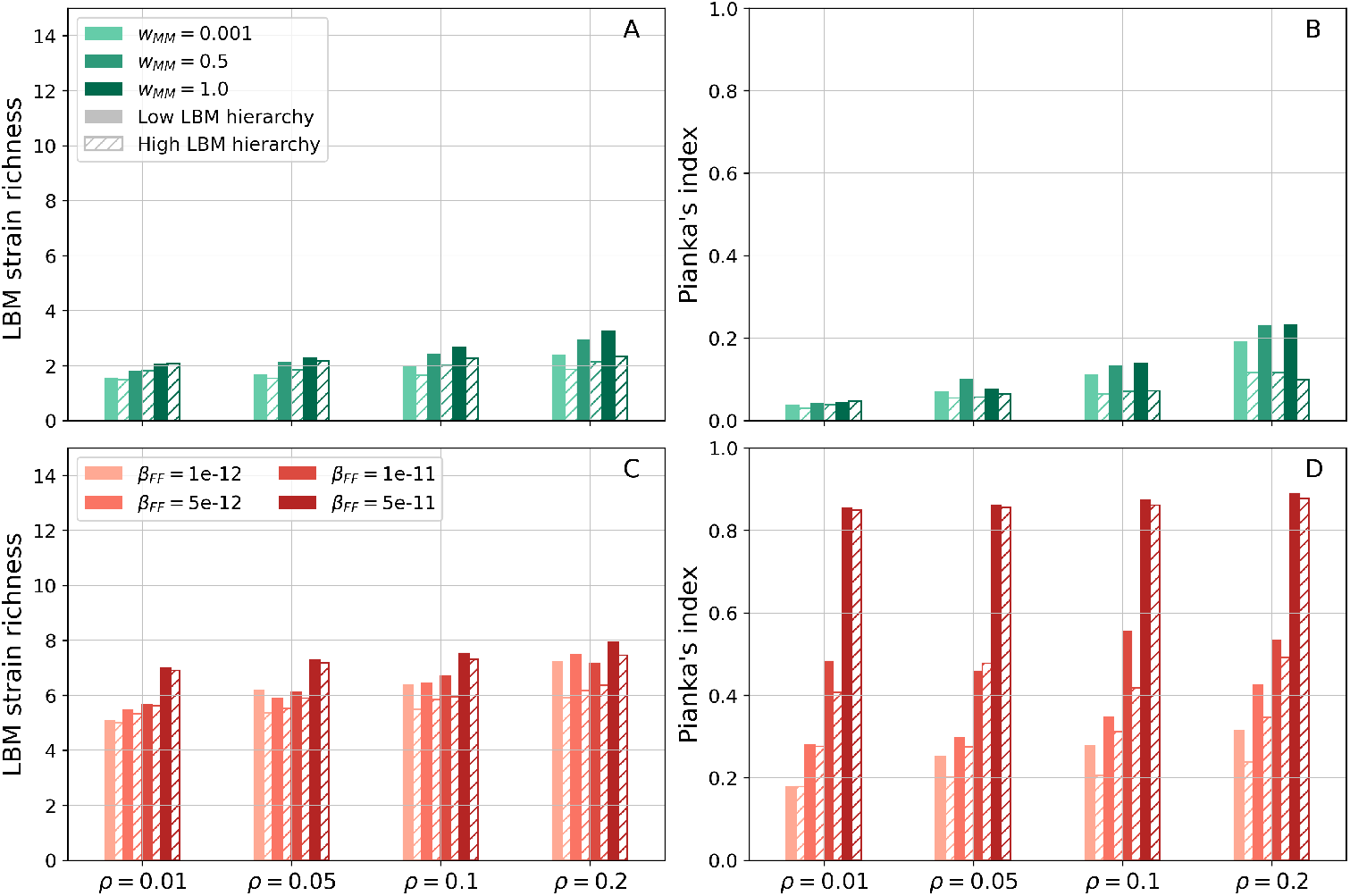
Viral mixing under complete cross-immunity. Results mirror panels B,C,E,F from Fig. 7 in the main manuscript, under the assumption of complete cross-immunity (*σ* = 0). Increasing cross-immunity lowers strain richness in any setting as individual strains face increased competition. Nonetheless, increasing cross-immunity does not significantly affect overlap between LBMs.

